# Unveiling the regulatory network controlling natural transformation in lactococci

**DOI:** 10.1101/2024.02.08.579460

**Authors:** Frédéric Toussaint, Marie Henry de Frahan, Félix Poncelet, Jean-Marc Ladrière, Philippe Horvath, Christophe Fremaux, Pascal Hols

## Abstract

*Lactococcus lactis* is a lactic acid bacterium of major importance for food fermentation and biotechnological applications. The ability to manipulate its genome quickly and easily through competence for DNA transformation would accelerate its general use as a platform for a variety of applications. Natural transformation in this species requires the activation of the master regulator ComX. However, the growth conditions that lead to spontaneous transformation, as well as the regulators that control ComX production, are unknown. Here, we identified the carbon source, nitrogen supply, and pH as key factors controlling competence development in this species. Notably, we showed that these conditions are sensed by three global regulators (i.e., CcpA, CodY, and CovR), which repress *comX* transcription directly. Furthermore, our systematic inactivation of known signaling systems suggests that classical pheromone-sensing regulators are not involved. Finally, we discovered that the ComX-degrading MecA-ClpCP machinery plays a predominant role based on the identification of a single amino-acid substitution in the adaptor protein MecA of a highly transformable strain. Contrasting with closely-related streptococci, the master competence regulator in *L. lactis* is regulated both proximally by general sensors and distantly by the Clp degradation machinery. This study not only highlights the diversity of regulatory networks for competence control in Gram-positive bacteria, but it also paves the way for the use of natural transformation as a tool to manipulate this biotechnologically important bacterium.

**IMPORTANCE:** Lactic acid bacteria (LAB) play important roles in our daily lives as members of our microbiota or as starters of dairy products. Understanding the natural horizontal gene transfer mechanisms that shape their genomes will allow us to better control and understand their evolution over time. The DNA transformation machinery is found in all beneficial LAB species. With the exception of streptococci, however, the conditions of its activation remain unknown. In this study, the physiological conditions that activate competence for DNA transformation in *Lactococcus lactis*, the most important lactococcal species, were identified. We also unveiled the guardians of the master competence regulator ComX. In this species, it is directly repressed by global carbon and nitrogen regulators (CcpA and CodY) as well as the general stress system CovRS. Additionally, it was discovered that the Clp machinery degrading ComX plays a dominant role in the strict control of competence activation. In Gram-positive cocci, the hierarchical organization of these regulators for controlling competence development in *L. lactis* is unprecedented.

## INTRODUCTION

*Lactococcus lactis* is one of the most studied species among lactic acid bacteria for both basic and applied research. Initially, the food industry has used its ability to produce lactic acid for centuries to preserve a wide variety of fermented foods (e.g., cheese, buttermilk, and sour cream) (56). In addition, *L. lactis* is now largely exploited as a safe production platform for recombinant proteins due to their easy production and downstream purification (39). Finally, this species is of high interest as a vector for therapeutics or vaccine antigens (1, 58). Because of all of these possible applications, *L. lactis* has become a powerful living tool of tremendous interest across the scientific community in just a few decades (56).

One of the commonalities shared by this large subset of applications is the ability to easily modify the lactococcal genome. Aside from commonly used techniques such as electroporation, conjugation, or transduction, the use of competence for DNA transformation is an appealing alternative. Natural transformation is a horizontal gene transfer process in which the bacterium is able to capture, internalize, and recombine a naked DNA fragment present in the extracellular environment (6). This process is thus of major interest for a variety of reasons. First, natural transformation does not necessitate the involvement of a third party as required for conjugation or transduction (i.e., another cell or a bacteriophage, respectively). Second, the donor DNA could simply be a linear fragment obtained by polymerase chain reaction and added to the culture medium. Third, natural transformation provides a quick and accurate genome editing tool (13).

Competence activation is typically divided into two distinct phases (i.e., early and late). During the early phase, a stimulus provokes the expression of the central regulator of competence. The latter, in association with the RNA polymerase, will then trigger the expression of the competence (*com*) regulon, which corresponds to the late phase (8, 22, 34). This regulon codes for all proteins responsible for DNA transformation, from DNA capture outside the cell to recombination into the genome (for recent reviews, see (13, 41)). While the late-phase development and associated mechanisms are well conserved across prokaryotes, stimuli and regulators responsible for the expression of the central regulator of competence are highly species-specific (for a review, see (34)).

In the closely-related streptococci, competence was shown to be controlled by a pheromone quorum-sensing system that directly activates the central competence regulator ComX (σ^X^) (22). In four out of the six groups of streptococci (i.e, salivarius, mutans, pyogenes, and suis), the intracellular sensor ComR of the Rgg family was identified, in combination with its secreted and re-internalized communication peptide XIP (<u>C</u>omX-Inducing <u>P</u>eptide, ComRS system), as the main trigger of competence activation (21, 22, 38). In the two other groups (i.e., mitis and anginosus), a different communication system is in charge of competence activation (34). This system is based on a phospho-relay that is activated by the secreted communication peptide CSP (<u>C</u>ompetence-<u>S</u>timulating <u>P</u>eptide, ComCDE system). The peptide is not re-internalized in this case but instead interacts with the extracellular part of the histidine kinase from the two-component system (TCS) ComDE (46). After a phosphorylation cascade, the response regulator ComE∼P triggers *comX* expression (37). In *Bacillus subtilis*, a similar mechanism was identified with the ComXAP system, in which the communication peptide ComX is secreted in the extracellular environment to activate the TCS ComAP (49). This leads to the production of ComS, a small intracellular peptide that disrupts the interaction between the central regulator of competence ComK and the adaptor protein MecA (9, 43, 47). This loss of interaction prevents ComK from being degraded by the ClpCP machinery, allowing its accumulation and expression of the late *com* regulon (for a review, see (29)).

So far, no such mechanisms of early-phase activation have been identified in *Lactococcus* species. However, a systematic analysis of all sequenced and complete genomes of *L. lactis* revealed that ∼25% of the strains have a complete and intact set of late *com* genes along with the *comX* gene, and that this relatively small proportion of strains is primarily isolated from plants (13). Interestingly, two strains containing the entire set of late *com* genes, *Lactococcus cremoris* KW2 and *L. lactis* KF147, were challenged for their ability to be transformed by exogenous DNA upon artificial ComX overproduction (11, 40). In both cases, the controlled induction of ComX allowed transformation events, demonstrating the capacity of *L. cremoris* and *L. lactis* to develop and carry out natural transformation (11, 40). Besides, spontaneous natural transformation has never been observed in those two species (11, 40). While an increase in *comX* expression along with some late *com* genes was reported under carbon starvation conditions in *L. lactis* KF147, transformation attempts with linear plasmid DNA failed (18). In *L. cremoris* KW2 containing the *comX*-overexpression plasmid in non-induced conditions, low spontaneous transformation events were observed in mutant strains lacking MecA, ClpC, or ClpP (11). However, this was attributed to a weak leakage of the inducible promoter and the cell’s inability to degrade ComX via the MecA-ClpCP machinery, resulting in ComX accumulation above the competence activation threshold (11).

In this work, we report the first observations of spontaneous competence in multiple *L. lactis* strains after optimization of DNA transformation conditions. Notably, we identified three global nutritional/stress regulators that repress *comX* transcription directly. Furthermore, we discovered a plant-derived *L. lactis* strain capable to transform at high level due to a single amino-acid substitution in the adaptor protein MecA of the Clp machinery. Together, these findings show that competence development in *L. lactis* is negatively controlled by (i) global regulators sensing nutritional and environmental conditions to modulate ComX abundance and (ii) the Clp machinery degrading ComX to limit its availability, both levels of regulation ensuring tight control of competence activation in this species.

## RESULTS

### Spontaneous transformation of *L. lactis* is strain-dependent

An *in silico* screening of an in-house database of ∼300 genomes of *L. lactis* allowed the identification of 18 strains (16 of plant origin) harboring the complete and supposedly intact set of *com* genes required for DNA transformation (*comX* and 16 late genes) (Table S1) (11). These 18 strains were initially tested for their potential ability to perform natural transformation. For this purpose, we used a plasmid-based system that allows *comX* overexpression by xylose induction (pGhP_xylT_*comX*) (11). This plasmid was successfully electroporated in 16 out of the 18 selected strains. Transformation assays were performed in M17GX (0.1% glucose-1% xylose) with a mutated version of the *rpsL* gene (*rpsL*\*, RpsL-K56R) as donor DNA, which confers streptomycin resistance (11). The results showed that 15 out of the 16 strains could be transformed at a rate between ∼10^-7^ and ∼10^-2^ (Fig. S1A), highlighting a high correlation between the presence of a complete set of competence genes and the functionality of the natural transformation machinery.

After this validation step, we challenged these strains for spontaneous transformation in rich medium M17G (0.5% glucose) with *rpsL*\* DNA. Notably, strain DGCC12653 of plant origin was the only one to show spontaneous transformation events in those conditions (Fig. 1A). We validated that natural transformation took place through the transformation machinery since a deficiency in ComX or in the DNA channel ComEC abolished competence (Fig. 1A). Moreover, a time-lapse experiment showed that transformation occurred at the entry of the stationary growth phase (Fig. 1B), contrasting with competence development in streptococci that takes place during the early logarithmic phase (8, 22).

**Figure 1.**
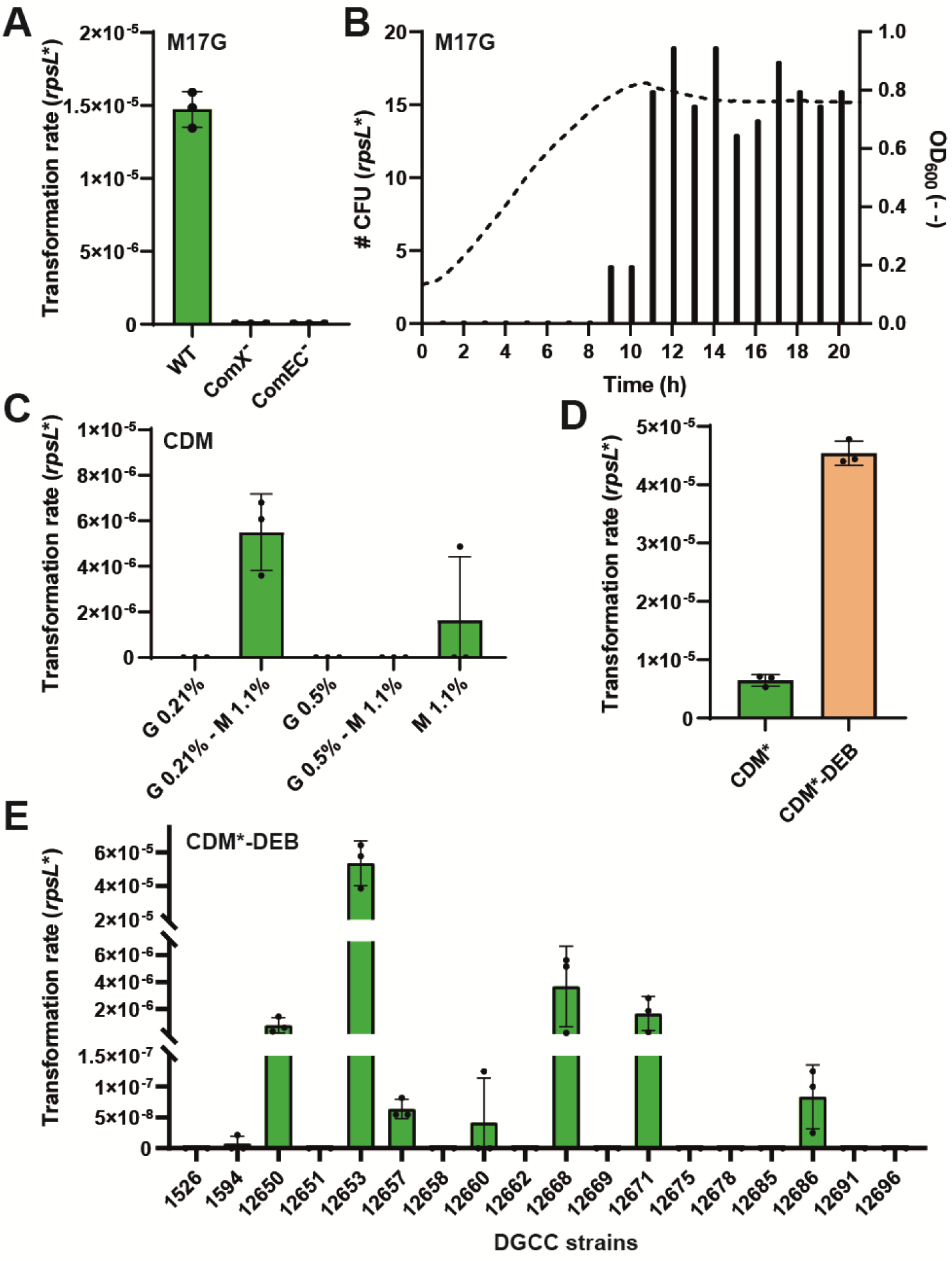
Effects of medium composition on spontaneous natural transformation. (A) Effects of the inactivation of ComX or ComEC on DGCC12653 transformability in M17G medium. (B) Timing of transformation events (bars, number of streptomycin-resistant CFUs) during cell growth (dotted line) in M17G medium. (C) Effects of glucose (G)-maltose (M) concentrations (%, w/v) on transformation rate in CDM. (D) Effects of aspartate, glutamate, and nitrogen bases [D, E, and B, respectively] omissions on transformation rate in CDM* (CDM Glu 0.21% - Mal 1.1%). The optimized competence-inducing medium was named CDM*-DEB. (E) Transformation rates of 18 *L. lactis* strains harboring the complete set of competence genes in CDM*-DEB. All transformation assays were performed with *rpsL*\* as donor DNA (20 µg ml^-1^), added at time zero. Cells were spread after ∼24 hours of culture (except in panel B). Dots show the values for biological triplicates (panel A) or technical triplicates (panels C, D, and E); mean values ± standard deviations.

Together, these results showed that spontaneous transformation of *L. lactis* is highly strain-dependent in regular rich medium and develops at the entry of the stationary growth phase.

### The carbon source and diauxic shift are triggers of spontaneous transformation

In order to explore the influence of various compounds in the growth medium on competence activation, we investigated transformation of strain DGCC12653 in chemically-defined medium (CDM) (53). As the carbon source was previously shown to have a strong impact on spontaneous transformation in many bacteria (2), we tested a large set of sugars using phenotype microarrays (Biolog plates). While no transformation was observed in CDM with glucose (0.5%) as the sole carbon source, we obtained transformants at low levels (∼10^-8^) with alternative sugars mainly originating from plants (i.e. maltose, xylose, cellobiose, galactose, and arabinose). As glucose starvation was previously reported to trigger the expression of competence genes in *L. lactis* (18, 48), we attempted to combine these two effects by testing the influence of a diauxic shift on transformability. Transformation assays were performed in CDM supplemented with glucose and maltose in different ratios (from 0 to 1%). Surface response of design of experiments (DoE) highlighted the most efficient mix between these two sugars (0.21% glucose-1.1% maltose) (Fig. 1C and Fig. S1B). This competence-inducing medium (named CDM*) allowed a reproducible transformation rate of ∼5 × 10^-6^ with *rpsL*\*. We also investigated the effect of amino acid and nitrogen base (purines and pyrimidines) omissions on spontaneous transformation. By removing aspartate, glutamate and nitrogen bases from the CDM medium (named CDM*-DEB), the transformation rate was ∼10-fold increased, reaching ∼4.5 × 10^-5^ (Fig. 1D). Finally, transformation assays (*rpsL*\* DNA) in this optimized medium were performed with the 18 *L. lactis* strains harboring the complete set of late *com* genes. In those conditions, spontaneous transformation was observed in 8 out of the 18 wild-type strains, with a transformation rate ranging from ∼10^-8^ to ∼10^-5^. Among these strains, DGCC12653 remained the most efficient spontaneous transformer (Fig. 1E).

Together, these results highlight that a sugar diauxic shift improves the transformation rate of a range of *L. lactis* strains, showing that growth on alternative carbon sources to glucose is a key physiological trait to activate spontaneous transformation in this species.

### The carbon catabolite regulator CcpA is a direct repressor of *comX*

To further investigate the role of the glucose-maltose diauxic shift on spontaneous competence of strain DGCC12653, we monitored the expression of the *comX* gene. For this purpose, a luminescence reporter system was designed by cloning the luciferase genes *luxAB* under the control of the *comX* promoter (P*_comX_*) in a low-copy number plasmid (pGhP_comX_*luxAB*). Luminescence assays showed that *comX* expression was boosted after seven hours of culture, which corresponds to the diauxic shift between glucose starvation and a second growth consuming maltose (Fig. 2A). Parallelly, we performed transformation assays by adding *rpsL*\* DNA at the beginning of the culture and monitored the transformation rate every hour after DNA addition. While no transformation was observed before seven hours of culture, transformation events occurred at the diauxic shift and increased until ten hours post DNA addition to reach a maximum transformation rate of ∼6 × 10^-5^ (Fig. 2A). This revealed that the diauxic shift stimulates ComX production at a sufficient level to allow competence activation.

**Figure 2.**
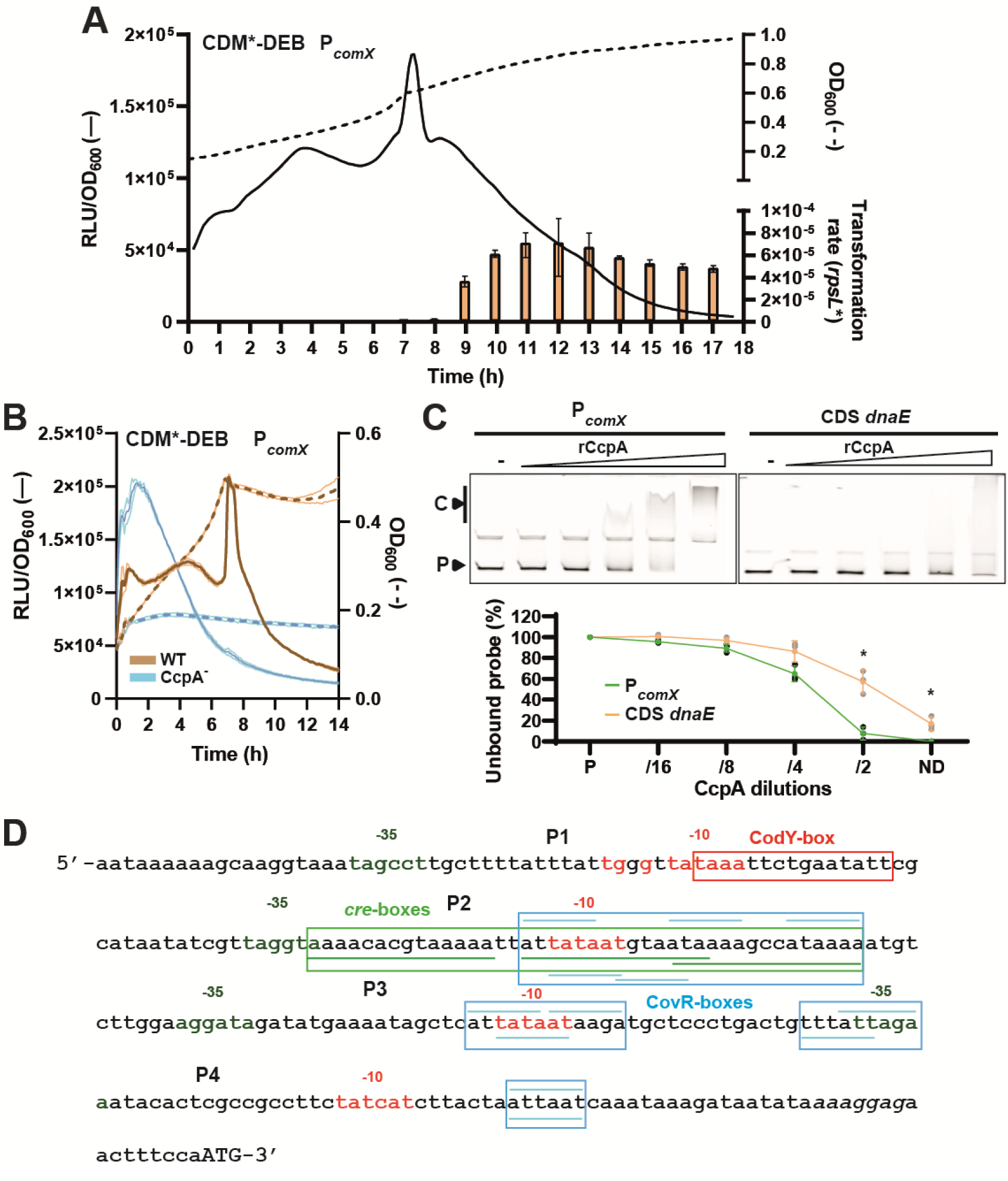
Involvement of CcpA in competence regulation. (A) Timing of transformation rate (orange bars) during cell growth (dotted line; OD_600_) and kinetics of P*_comX_* specific luciferase activity (solid line; RLU/OD_600_) in CDM*-DEB. Bars show mean values of technical triplicates ± standard deviations. Transformation assays were performed as reported in figure 1. (B) Growth (dotted lines; OD_600_) and kinetics of P*_comX_* specific luciferase activity (solid lines; RLU/OD_600_) monitored over time for DGCC12653 WT (orange) and CcpA^-^ strains (blue). Solid and dotted dark lines are representative of the mean of technical and biological triplicates (WT and CcpA^-^, respectively), and light lines are representative of standard deviation. (C) EMSAs (top panels) performed with a gradient of purified CcpA (rCcpA) on P*_comX_* (left) and a 150-bp fragment of the CDS of *dnaE* as negative control (right). Lanes without rCcpA are indicated by a minus sign. C and P indicate the rCcpA-DNA complex and the unbound probe, respectively. Gels displayed are representative of technical triplicates. The percentage of unbound probe (bottom panel) for P*_comX_* (green line) and CDS *dnaE* (orange line) was calculated using the Amersham Typhoon analysis software. ND and P stand for non-diluted rCcpA and probe-alone condition, respectively. Statistical *t* test was performed for each dilution in comparison to the negative control (*n*=3; *, *P* < 0.05). (D) Mapping of regulation elements identified in P*_comX_* (intergenic region). CodY-, *cre*- and CovR-boxes are surrounded in red, green, and blue, respectively. Top and bottom blue lines correspond to CovR boxes identified on the top and bottom strands, respectively. Putative −35 and −10 boxes of mapped vegetative promoters (P1 to P4) are colored in red and dark green, respectively. The Shine-Dalgarno (italics) and start codon (capitals) of *comX* are mapped at the 3’end.

As we observed a stimulating effect of the diauxic shift on competence development, we questioned the role of the global transcriptional regulator CcpA (carbon catabolite control protein A) in this phenomenon. CcpA is involved in carbon catabolite repression (CCR) in Gram-positive bacteria and plays a role in the diauxic shift between the preferred glucose catabolism and alternative sugar utilization pathways in the closely-related species *L. cremoris* (54). A *ccpA-*deleted mutant of strain DGCC12653 was constructed and transformed with the P*_comX_*-*luxAB* reporter system. We observed an increased expression of *comX* in the early growth phase (∼2-fold) along with a major growth defect for the *ccpA* mutant compared to the wild-type in optimized conditions (Fig. 2B). We also performed transformation assays in the same conditions that showed a decreased transformation rate when *ccpA* was deleted (Fig. S2A). Although *comX* expression was slightly unleashed in the *ccpA* mutant, the constitutive CcpA deficiency has a global impact on growth and carbon metabolism that could be incompatible with the transformation process (e.g., alterations of cell-wall, ATP availability, or cell cycle).

To test whether CcpA is directly involved in competence regulation by binding P*_comX_*, we performed electrophoretic mobility shift assays (EMSAs) with purified CcpA of strain DGCC12653 (Fig. S2B). The purified protein (CcpA with N-terminal 6-His tag; named rCcpA) was incubated with fluorescent Cy3-labelled P*_comX_* (^Cy3^P*_comX_*) as a DNA probe (Fig. 2C), which corresponds to the intergenic region located upstream of the *comX* gene. The results showed that rCcpA significantly interacts with P*_comX_* compared to the control probe (CDS of *dnaE*) (Fig. 2C). Thanks to *in silico* analyses and different DNA probes (Fig. S2C-D), we were able to identify three potential CcpA-binding sites (*cre*-box consensus, WGWAARCGYTWWMA) (69) that overlap the putative vegetative promoter P2 in the upstream intergenic region of *comX* (Fig. 2D).

Together, these results show that a sugar diauxic shift activates *comX* expression at the transcriptional level. They also suggest that CcpA directly represses P*_comX_* in glucose conditions, a repression effect that is potentially released on alternative carbon sources such as maltose.

### The nitrogen global regulator CodY is a direct repressor of *comX*

During the optimization process of CDM composition, we observed that branched-chain amino acids (BCAAs; Leu, Val, and Ile) are required to sustain the growth of strain DGCC12653 but that an excess of isoleucine decreases *comX* expression and spontaneous transformation in optimized conditions (Fig. 3A and B). As BCCAs are well known to control the activity of the nitrogen transcriptional regulator CodY in *L. lactis* and *L. cremoris* (12, 26, 27, 69), we investigated its potential role in the control of competence. For this purpose, we generated a *codY*-deleted mutant of strain DGCC12653 that was transformed with the P*_comX_*-*luxAB* reporter system. CodY inactivation resulted in a nearly 10-fold increase in *comX* expression at the diauxic shift and a drastic improvement of the transformation rate (maximum of ∼ 10^-3^) compared to the wild type in optimized conditions (Fig. 3C-E). The activation of the late *com* phase in the *codY* mutant was also validated using a P*_comGA_*-*luxAB* reporter system (Fig. 3D). In addition, a RNAseq analysis was performed to compare the wild type and the *codY* mutant using total RNAs extracted during the diauxic shift (Table S1). This analysis confirmed the higher expression of *comX* along with the complete late *com* regulon in the CodY-deficient strain. These results suggest that CodY is somehow repressing competence development by downregulating the expression of *comX* and late *com* genes.

**Figure 3.**
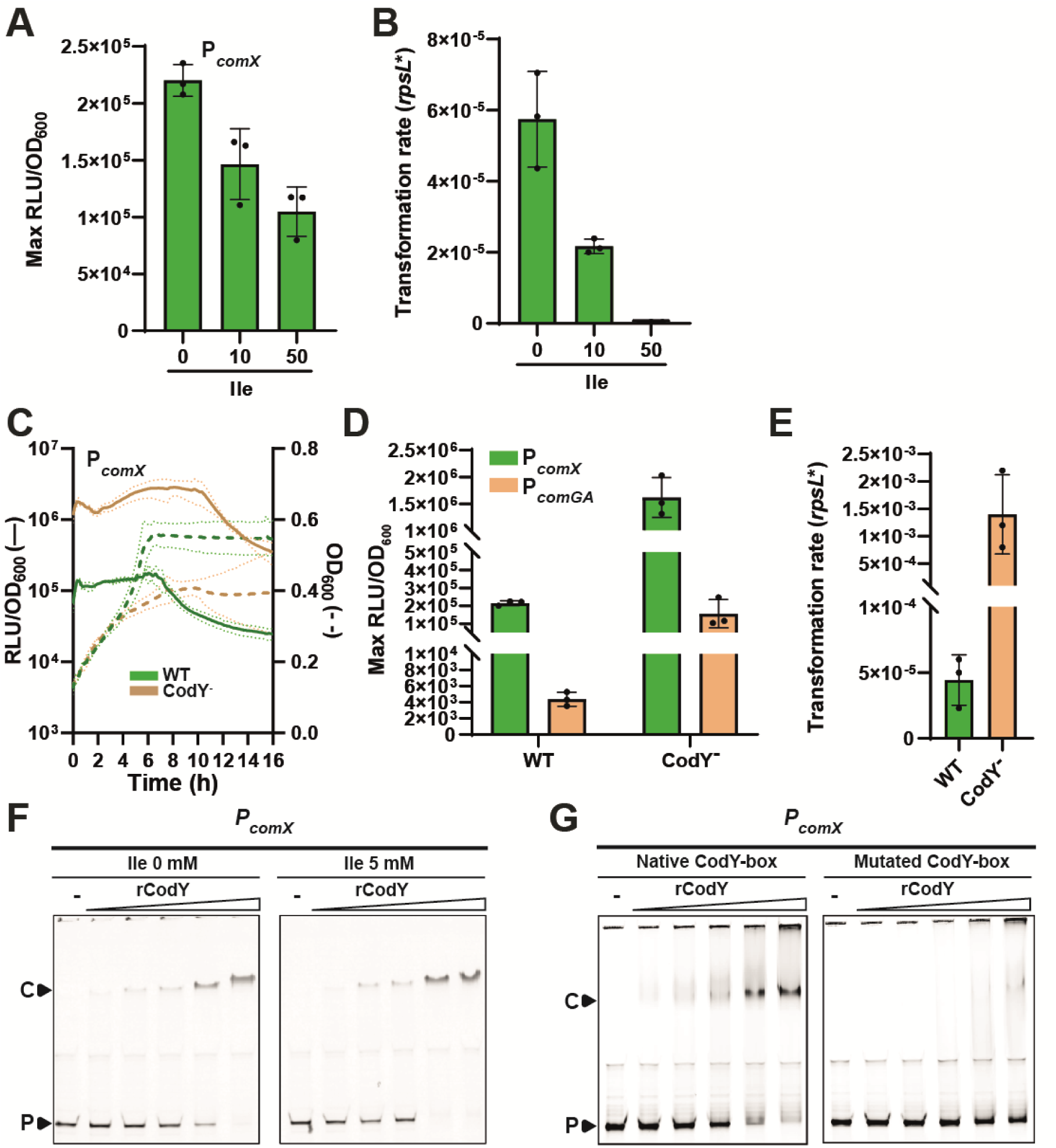
Involvement of CodY in competence regulation. (A) Effect of isoleucine excess on P*_comX_* activity in DGCC12653 WT. Data show maximum specific luciferase activity (Max RLU/OD_600_) observed at the diauxic shift for different concentrations of isoleucine (Ile; 0, 10, and 50 mM) added to the initial concentration (0.76 mM) in CDM*-DEB. (B) Effect of isoleucine excess (conditions as in panel A) on transformability of DGCC12653 WT after overnight culture with donor DNA (C) Growth (dotted lines; OD_600_) and kinetics of P*_comX_* specific luciferase activity (solid lines, RLU/OD_600_) monitored over time for WT (green) and CodY^-^ strains (brown) in CDM*-DEB. Solid and dotted dark lines are representative of the mean of technical and biological triplicates (WT and CodY^-^, respectively), and light lines are representative of standard deviation. (D) Effect of *codY* deletion on activation of early (P*_comX_*) and late (P*_comGA_*) competence phases. Data show maximum specific luciferase activity (Max RLU/OD_600_) observed at the diauxic shift using either P*_comX_*-*luxAB* (green) or P*_comGA_*-*luxAB* (light brown) reporter fusions in CDM*-DEB. (E) Effect of *codY* deletion on transformability. Data show transformation rates observed for WT (green) and CodY^-^ strains (light brown) in CDM*-DEB after overnight culture with donor DNA. Dots show the values for biological triplicates (CodY^-^) or technical triplicates (WT), mean values ± standard deviations. (F) Effect of isoleucine on CodY binding to P*_comX_.* EMSAs performed with a gradient of purified CodY (rCodY) on P*_comX_* without (left) or with 5 mM isoleucine (right). (G) Identification of the CodY-box in P*_comX_*. EMSAs comparing rCodY binding to native (left) and mutated (right) CodY-boxes. In the mutated CodY-box, the two central nucleotides CG were swapped with AA. Lanes without rCodY are indicated by a minus sign. C and P indicate the rCodY-DNA complex and the unbound probe, respectively.

To question the possible direct interaction between CodY and P*_comX_*, we purified CodY of strain DGCC12653 (CodY with N-terminal 6-His tag; named rCodY) for binding assays (Fig. S3A). EMSAs performed with ^Cy3^P*_comX_* confirmed the direct binding of rCodY to P*_comX_*. Subsequent analyses revealed that rCodY binding was favored by the presence of BCAAs with the strongest effect observed with isoleucine (5 mM), but not by GTP as previously reported in *L. lactis* (26) (Fig. 3F and Fig. S3B). We also identified a CodY-binding motif (CodY-box T<u>A</u>AA<u>TTC</u> T <u>GAA</u>T<u>ATT</u>, conserved positions underlined (12)) that overlaps the −10 box of the putative vegetative promoter P1 in the upstream intergenic region of *comX* (Fig. 2D). To confirm the CodY-box, we mutated the two central nucleotides CG into two AA (T<u>A</u>AA<u>TT</u>A T A<u>AA</u>T<u>ATT</u>), and showed that rCodY binding was drastically reduced (Fig. 3G). In addition, the mapping of RNAseq reads on P*_comX_* displays a strong increase of transcripts around the CodY box for the CodY-deficient strain compared to the wild type. However, most of those reads result from a transcriptional readthrough from the upstream *ezrA* gene (Fig. S3C).

These results show that CodY represses competence development by downregulating *comX* expression through a direct binding on P*_comX_* in *L. lactis*. This highlights that the central competence gene *comX* of this species is regulated by the cell nutritional status through the sensing of both carbon and nitrogen growth substrates.

### The environmental stress sensor CovRS is a direct repressor of *comX*

In order to determine if competence in *L. lactis* is controlled by a cell-to-cell communication system or stress sensor(s), as reported in other Gram-positive bacteria (8, 22, 34), we systematically inactivated all members of the Rgg family and TCSs. Ten Rgg and eight complete TCS (response regulator [RR] and histidine kinase [HK] in tandem) encoding genes were identified in the genome of strain DGCC12653 (Table S2). All these genes were successfully inactivated by deletion, except the VicRK (WalRK) system that was previously reported to be essential in Gram-positive bacteria (57). The collection of 17 mutants was tested for *comX* expression (P*_comX_*-*luxAB*) and spontaneous transformation in the optimized medium. All ten *rgg* mutants behaved as the wild-type strain, with less than a two-fold impact on transformation and luminescence (Fig. S4A and B). Regarding the seven TCS mutants (tandem inactivation of RR/HK), similar analyses revealed that CovRS (DGCC12653_01910-01915) inactivation allowed both an increase in *comX* expression (up to 15-fold) and transformation rate (up to ∼ 10^-3^), while all other deletion mutants did not significantly differ from the wild-type strain (Fig. 4A-C, and Fig. S4C and D). A RNAseq analysis was also performed to compare the transcriptomes of the *covRS* mutant with the wild type. As the CovRS-deficient strain displayed a strong growth defect in CDM*-DEB, both the wild type and *covRS* mutant were propagated in CDM*. Total RNAs were extracted from cells collected during the diauxic shift, when spontaneous transformation was previously shown to take place. This experiment confirmed the increase of *comX* expression, as well as the complete set of late *com* genes, in the *covRS* mutant compared to the wild type (Table S1). In addition, the P*_comGA_*-*luxAB* reporter system of the late competence phase was strongly induced (Fig. 4B), confirming RNAseq data and high transformation rate. As CovRS was previously shown to respond to pH in streptococci (35, 45, 50), we monitored the transformation rate in a range of initial pH values (pH 5.5 to 8.0). While spontaneous transformation in the wild-type strain was abolished below pH 6.6, the *covRS* mutant remained transformable in a much larger range of low pH values (Fig. 4D). In addition to the nutritional sensors CcpA and CodY, these results show that the stress sensor CovRS represses *comX* expression and thus transformation efficiency in *L. lactis* DGCC12653.

**Figure 4.**
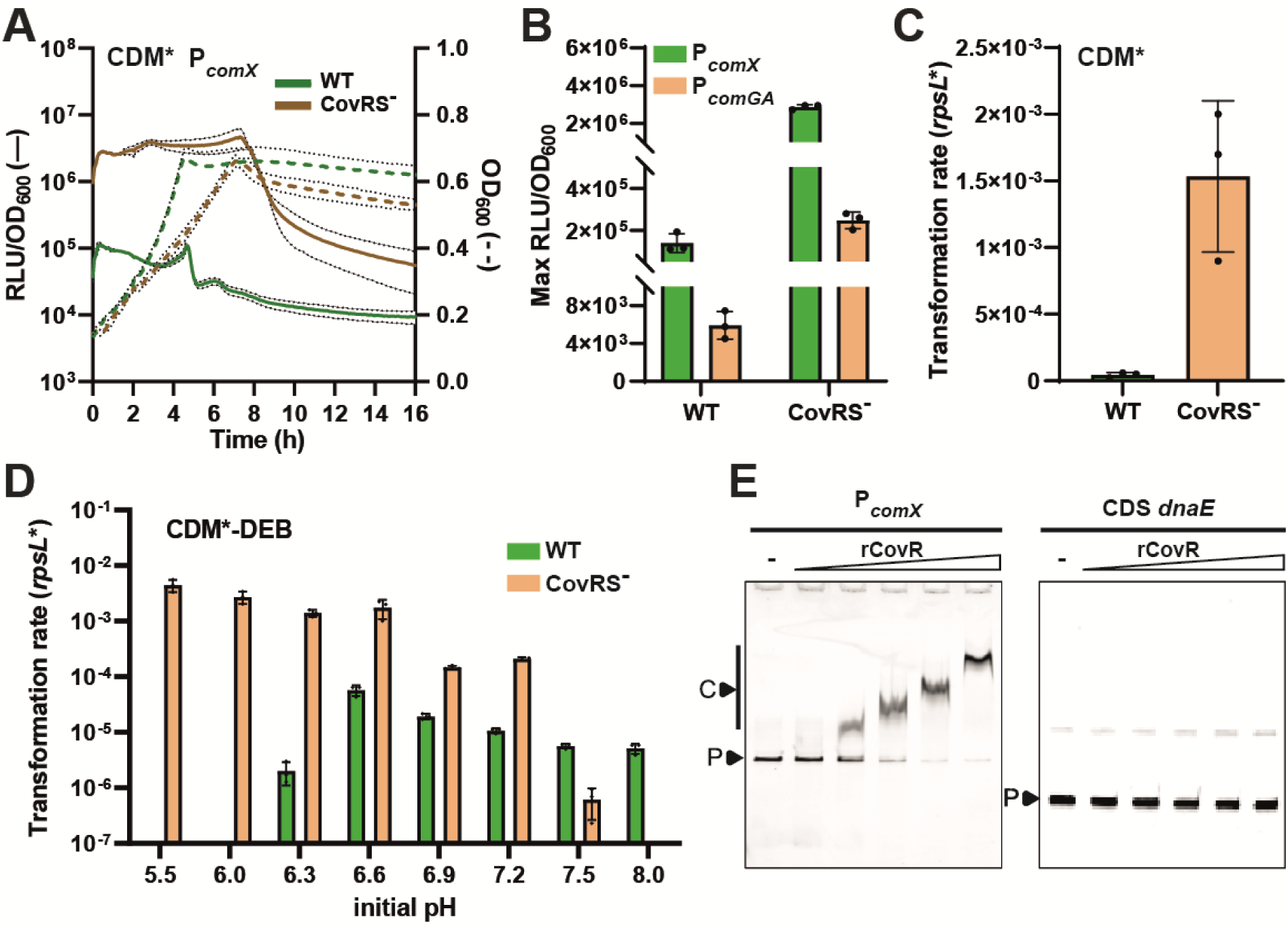
Involvement of CovRS in competence regulation. (A) Growth (dotted lines, OD_600_) and kinetics of P*_comX_* specific luciferase activity (solid lines, RLU/OD_600_) monitored over time for DGCC12653 WT (green) and CovRS^-^ strains (brown) in CDM*. Solid and dotted dark lines are representative of the mean of technical and biological triplicates (WT and CovRS^-^, respectively), and light lines are representative of standard deviation. (B) Effect of *covRS* deletion on activation of early (P*_comX_*) and late (P*_comGA_*) competence phases. Data show maximum specific luciferase activity (Max RLU/OD_600_) observed at the diauxic shift using either P*_comX_*-*luxAB* (green) or P*_comGA_*-*luxAB* reporter fusions in CDM*. (C) Effect of *covRS* deletion on transformability. Data show transformation rates observed for WT (green) and CovRS^-^ (light brown) strains in CDM* after overnight culture with donor DNA. (D) Effect of the initial pH on transformability. Data show transformation rates observed for WT (green) and CovRS^-^ strains (light brown) after overnight culture with donor DNA in CDM*-DEB buffered at different initial pHs. Dots show the values for biological triplicates (CovRS^-^) or technical triplicates (WT), mean values ± standard deviations. (E) EMSAs performed with a gradient of purified CovR (rCovR) on P*_comX_* (left) and the CDS of *dnaE* as negative control (right). Lanes without rCovR are indicated by a minus sign. C and P indicate the rCovR-DNA complex(es) and the unbound probe, respectively. Gels displayed are representative of technical triplicates.

To question the possibility that CovR directly regulates competence by binding P*_comX_*, we purified CovR from strain DGCC12653 (CovR with N-terminal 6-His tag; named rCovR) (Fig. S5A) and performed EMSAs with P*_comX_* as a probe. Interestingly, EMSAs showed that *in vitro*-phosphorylated CovR was able to efficiently bind the full-length intergenic region (Fig. 4E), thus acting as a direct repressor of *comX* expression in *L. lactis*. EMSAs assays with different DNA probes allowed us to identify multiple CovR-binding regions covering putative promoters P2, P3, and P4 (Fig. S5B-D and Fig. 2D). *In silico* analysis of these regions revealed multiple CovR-boxes (ATTARA) that were previously shown to be CovR-binding sites in streptococci (Fig. 2D) (7, 19). We also showed that the mapping of RNAseq reads on P*_comX_* displays a strong increase of transcripts downstream of the promoter P2 for the CovRS-deficient strain compared to the wild type, corroborating EMSA results (Fig. S5E).

Altogether, these results indicate that classical cell-to-cell communication systems (Rgg and TCS) reported for competence control in Gram-positive bacteria (22) are not involved in *comX* activation in *L. lactis*. Conversely, these results confirm that *comX* expression is under the direct control of multiple global regulators, including the general stress sensor system CovRS, which responds to a range of environmental stimuli in closely-related species (23, 25, 35, 45, 50).

### The Clp degradation machinery plays a central role in competence repression

In order to extend the role of global regulators in the control of natural transformation in other *L. lactis* strains, we generated *codY* and *covRS* mutants in three additional *L. lactis* strains (DGCC12651, DGCC12671 and DGCC12678) whose natural transformation was validated by the artificial *comX* overexpression system (Fig. S1A). Surprisingly, while *comX* expression was similarly increased in those mutants compared to strain DGCC12653 (Fig. S6A), none of these mutant strains (neither *codY* nor *covRS* deleted mutant) showed spontaneous transformation with *rpsL*\* DNA in optimized conditions. These results suggest an additional layer of regulation at the post-transcriptional level between ComX production and transcriptional activation of the late *com* genes.

As we previously reported that the ComX-degrading MecA-ClpCP machinery negatively affects artificially-induced natural transformation of *L. cremoris* (11), we compared the protein sequences of MecA, ClpC, and ClpP of these three strains with those from strain DGCC12653. While alignments of ClpC and ClpP proteins showed no specific amino acid variation, the adaptor protein MecA displayed a single amino acid change (L125R) that is only present in strain DGCC12653 (Fig. S6B). To test whether the L125R substitution could be responsible for the spontaneous transformability of DGCC12653, we reciprocally swapped *mecA* genes between strains DGCC12653 (MecA-R125; rare) and DGCC12671 (MecA-L125; common) (Fig. 5A). We also generated *mecA* deletion mutants (MecA^-^) of both genetic backgrounds. Notably, we observed that DGCC12653 that produces the common version of MecA (L125) lost its capacity to transform exogenous DNA. Conversely, the production of the rare version of MecA (R125) in a non-spontaneously transformable strain (DGCC12671) activated natural transformation at a rate comparable to a MecA^-^ strain (Fig. 5B). In addition, we were able to delete *mecA* in 12 out of the 18 *L. lactis* strains initially selected. Natural transformation assays performed in four different competence-inducing media showed that *mecA* deletion allowed transformation events to different extents in eight out of the 12 MecA^-^ strains (Fig. 5C).

**Figure 5.**
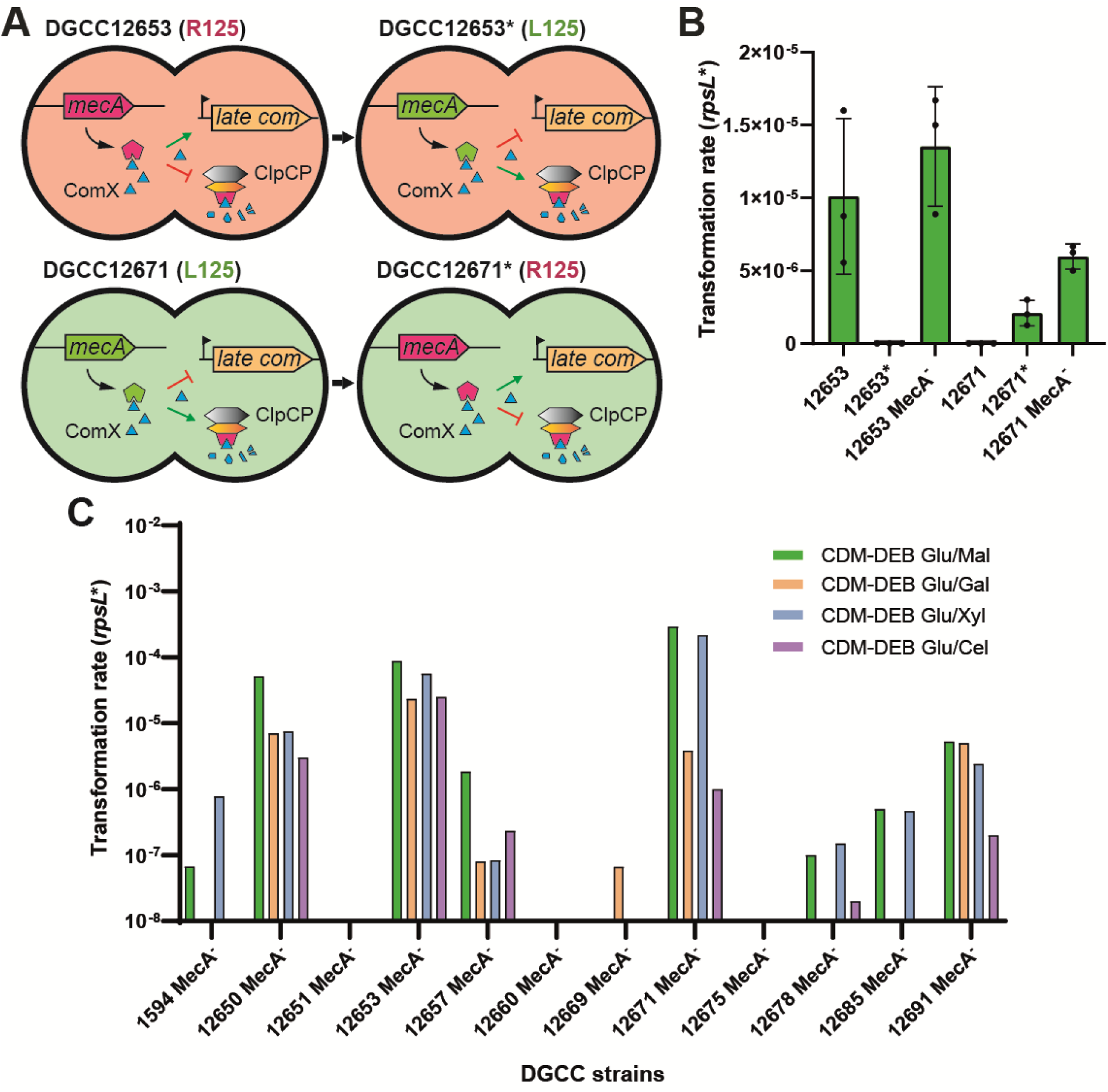
Involvement of MecA in competence regulation. (A) Schematic view of the swap of *mecA* genes between strains DGCC12653 (salmon) and DGCC12671 (light green). Genes encoding MecA-R125 (rare) and MecA-L125 (common) are colored in pink and green, respectively. Green and red lines illustrate activated or repressed pathways, respectively. Black lines display gene expression. (B) Effects of MecA-L125R and -R125L substitutions on transformability. Data show transformation rates observed for DGCC12653 WT (MecA-R125), DGCC12653* (MecA-L125), DGCC12671 WT (MecA-L125), DGCC12671* (MecA-R125), and MecA^-^ (*mecA* deletion) strains after overnight culture with donor DNA in CDM*-DEB. Dots show the values for biological triplicates (mutant strains) or technical triplicates (WT), mean values ± standard deviations. (C) Effects of *mecA* deletion (MecA^-^) on spontaneous transformability in 12 *L. lactis* strains. Data show transformation rates in four different media composed of CDM-DEB with glucose (Glu) 0.21% (w/v) and a second plant sugar at 1.1% (w/v) (maltose [Mal], green; galactose [Gal], light brown; xylose [Xyl], blue; and cellobiose [Cel], purple).

Together, these results showed that MecA plays a crucial role in competence repression and that the single substitution MecA-L125R explains the unique transformation behavior of strain DGCC12653. Contrasting with the low impact of the Clp degradation machinery on competence control in streptococci (22, 59, 62), this machinery plays a dominant role at the global level of competence regulation in *L. lactis*.

## DISCUSSION

In this work, we highlighted a complete rewiring of the regulatory network controlling competence in *L. lactis* when compared to closely-related streptococci (Fig. 6). Although ComX is the common central regulator of competence in both, the proximal regulation of its activation does not appear to be controlled by a cell-to-cell communication module like in streptococci, but rather by global regulators sensing cellular nutritional status (CcpA and CodY) or stress conditions (CovRS) (Fig. 6A). By comparison, global regulators are distantly controlling ComX by modulating the communication module in streptococci (for a review, see (22)) (Fig. 6B). Moreover, the post-transcriptional layer of regulation through ComX degradation by the MecA-ClpCP machinery plays a predominant role in *L. lactis*, while it is more seen as a secondary locking device in streptococci (3, 14, 22, 55, 59, 62). In addition, the kinetics of competence activation during growth on a single carbon source are totally different: stationary phase in *L. lactis versus* transitory during early logarithmic growth for streptococci, controlled by the on/off switch of the communication module (8, 22). Notably, the timing of competence activation and the network hierarchy share some similarities with competence regulation in bacilli, where the central transcriptional regulator ComK is proximally modulated by global regulators and primarily controlled through degradation, which may, or may not, be regulated by a cell-to-cell signaling system. (Fig. 6B) (29, 33)).

**Figure 6.**
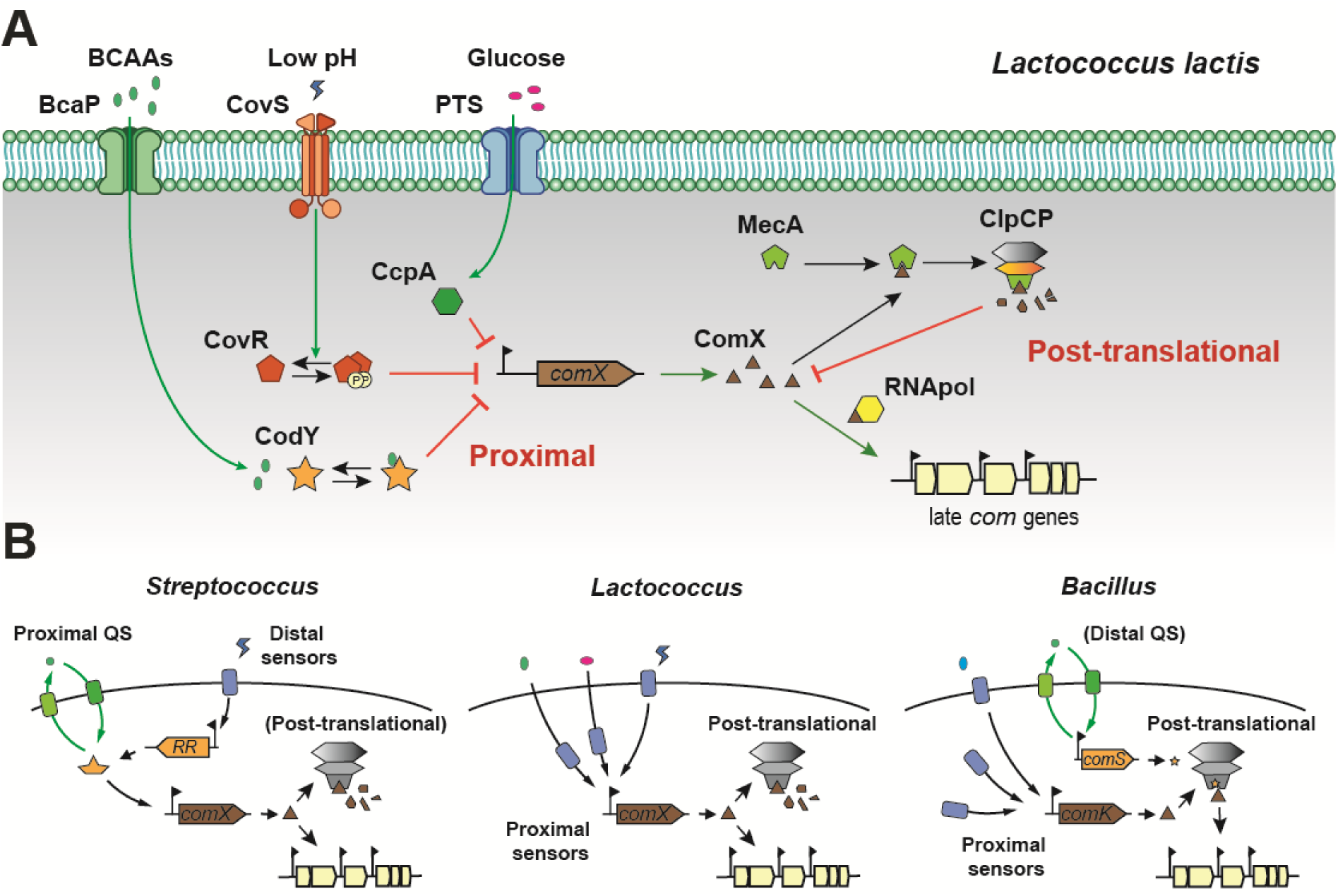
Competence control integrating proximal-distal transcriptional and post-translational regulation of the central regulator in *Lactococcus lactis* and closely-related species. (A) Model of competence regulation in *L. lactis*. The transcription of *comX* (proximal regulation) is repressed by three regulators, sensing either the cellular nutritional status (CcpA and CodY, green and orange, respectively) or stress conditions (CovRS, red). When environmental/stress conditions are appropriate, ComX (brown) production is unrepressed. Then, ComX can interact with the adaptor protein MecA (light green) for its degradation by the ClpCP machinery (light orange and gray, respectively), keeping ComX concentration at a low level (post-translational regulation). When ComX abundance overcomes the degradation machinery (probably in a subpopulation), it can interact with the RNA polymerase (yellow) to trigger the expression of late *com* genes and, in turn, natural transformation. (B) Comparison of competence regulation at the global level between *Lactococcus*, *Streptococcus*, and *Bacillus*. In *Streptococcus* (left panel), the proximal regulation of *comX* is performed by a quorum sensing (QS) system (ComRS or ComCDE, green). Additionally, distal regulation systems (e.g., TCSs, blue) responding to environmental cues are modulating the expression of the QS response regulator/sensor instead of *comX* as found in *L. lactis*. Finally, the post-translational regulation system (grey) plays a minor role in ComX degradation. In *Bacillus* (right panel), the proximal regulation of ComK is achieved by a range of transcriptional regulators (e.g., CodY, DegU, Rok, and AbrB). However, a distal regulation involving a QS system (ComXAP) triggers the production of ComS to inhibit ComK degradation in *B. subtilis*. In *Lactococcus* (middle panel), no QS system has been identified so far and proximal regulation of *comX* seems to be exclusively influenced by environmental conditions. Moreover, post-translational regulation was shown to be dominant in competence repression.

Among the competence triggers in *L. lactis*, we identified carbon nutritional status as a key signal. While spontaneous transformation in rich medium with glucose is activated at the entry of the stationary phase, it is abolished in CDM-glucose conditions (Fig. 1B and C). However, growth on alternative sugars mostly found in plants reactivates spontaneous competence at a low level in those conditions (Fig. 1C). This makes sense as strain DGCC12653 is a maize isolate with a high potential to catabolize alternative non-PTS (phosphoenolpyruvate:carbohydrate phosphotransferase system) sugars such as maltose, xylose, and arabinose (5). This also recalls the activation of competence by non-PTS sugars in Gram-negative bacteria (51). For instance, competence in *Vibrio cholerae* is induced by chitin oligomers but repressed by PTS-sugars such as glucose through CCR via the cAMP-activated CRP protein (2). In Gram-positive bacteria, the impact of sugars and the implication of CcpA-mediated CCR on competence development have not been deeply studied. Notably, our results showed that a diauxic shift between glucose and the non-PTS sugar maltose activates *comX* expression and transformability at the growth shift between the two sugars (Fig. 2A). They also showed that glucose starvation and the presence of maltose are both required to maximize spontaneous transformation (Fig. 1C). This corroborates previous observations showing that glucose starvation alone was unable to activate natural transformation in *L. lactis*, although a range of competence genes (including *comX*) were upregulated (18, 48). In addition, growth on maltose could be strongly delayed after glucose starvation from one experiment to the other without major effect on transformability. We hypothesize that the second sugar catabolism is required to maintain the energy demand needed for natural transformation (66). Interestingly, fluorescence microscopy analyses (fusion P*_comX_*-*gfp^sf^*) revealed a bimodality in competence activation at the diauxic shift since only a subpopulation of cells expressed *comX* at detectable levels (Fig. S7). This suggests a bet-hedging strategy for cells that are developing alternative physiological adaptations, as previously reported for diauxic growth with *L. cremoris* (54) or during competence activation in *B. subtilis* (36). In *L. cremoris*, CcpA-mediated CCR was previously shown to be involved in the diauxic shift due to glucose exhaustion (54). The inactivation of CcpA in strain DGCC12653 drastically alters growth and the dynamics of *comX* expression, with a two-fold upregulation at an early growth stage compared to the wild-type (Fig. 2B), potentially due to an absence of CcpA binding to the *comX* promoter. Similarly, a two-fold increase in *comX* expression was observed in a CcpA-negative strain of *Streptococcus oligofermentans* (63). In this case, CcpA was shown to indirectly control *comX* due to a post-transcriptional effect on the expression of the communication system ComCDE (63). However, CcpA inactivation in strain DGCC12653 leads to reduced transformability, as previously reported in different streptococci (60, 67, 68). CcpA is a pleiotropic regulator that controls, among others, the central carbon metabolism and a range of cell envelope components in lactococci and streptococci (4, 68, 69), whose constitutive inactivation could negatively alter the transformation process independently of an upshift in *comX* expression.

Besides the impact of the carbon source, the nitrogen status sensed by the global regulator CodY has a major impact on competence development in *L. lactis*. In this species, CodY is a global regulator involved in nitrogen metabolism (e.g., peptide and amino acid uptake, peptide degradation, *de novo* biosynthesis of amino acids, purine biosynthesis) that senses the intracellular pool of BCAA (mostly isoleucine) (12, 26, 27). We showed that an excess of isoleucine in the growth medium repressed *comX* expression and spontaneous transformation (Fig. 3A and B). We also identify that a lack of aspartate and glutamate in the growth medium improves transformability. As it was previously reported that both amino acids could be catabolized for their conversion into BCAA (16), we hypothesized that their removal could reduce the intracellular pool of BCAA and consequently CodY-mediated repression. In addition, CodY inactivation has strong effects, with a ∼10-fold higher activation of *comX*, a higher proportion of fluorescent cells post-diauxic shift, and a ∼100-fold increase in transformability (Fig. 3C-E and Fig. S7). This recalls the impact of CodY on the direct transcriptional repression of *comK* (52) and the importance of the amino-acid balance on transformability in *B. subtilis* (64). Finally, we showed that CodY directly binds the *comX* promoter, a region that was also identified by CHIPseq experiments as bound by CodY in *L. lactis* but without being further investigated (65). Although previously suggested in *L. lactis* (18), these results demonstrate that nitrogen sensing through CodY is central for the metabolic control of competence in this species.

In our search for competence-related quorum-sensing systems, we first scanned the genome for orthologous systems known in other Gram-positive species. As no such system was found, we systematically knocked out every Rgg (ten genes) and TCS (eight pairs of genes) and identified CovRS (TcsA in *L. cremoris*) as a major player in competence regulation in *L. lactis*. As found for CodY inactivation, its deletion dramatically increases *comX* expression, the proportion of fluorescent cells post-diauxic shift, and natural transformation (Fig. 4A-C and Fig. S7). We also showed that CovRS directly controls *comX* transcription in *L. lactis*, as confirmed by EMSA and RNAseq experiments (Fig. 4E and Fig. S5). Interestingly, we recently showed that this system indirectly controls *comX* expression in salivarius streptococci by modulating the ComRS signaling system via *comR* repression (35). CovRS is a well-known regulatory system of virulence genes in pathogenic streptococci, responding to various stimuli (e.g., [Mg^++^], defensins, osmotic stress, pH) (10, 23, 25, 45, 50). In *L. cremoris*, CovR (LlrA) was shown to be involved in acid stress resistance through arginine catabolism (42). Interestingly, we showed that competence sensitivity to low pH was abolished in the CovRS-deficient strain (Fig. 4D), corroborating its involvement in acid stress resistance. As CovRS and CodY are the dominant repressors of competence in *L. lactis*, we also generated a double *covRS codY* mutant to evaluate additive or synergic effects. Although transformability and maximum *comX* expression were not increased, *comX* expression became constitutive during growth (Fig. S8). This suggests that these two regulatory systems are controlling *comX* with a different temporality, synergistically during early logarithmic growth and then dominantly by CovRS until the entry into the stationary growth phase (Fig. S8). Besides CovRS, no other TCS or Rgg is involved in the repression or activation of the *comX* gene in strain DGCC12653 (Fig. S4A-D). As all identified quorum-sensing systems regulating competence in Gram-positive bacteria belong to those two families (8, 22, 34), this questioned the presence of a cell-to-cell communication system for the control of *comX* expression in *L. lactis*. A similar situation was observed in *Bacillus licheniformis*, where competence regulation does not involve the ComXAP-like quorum-sensing system found in *B. subtilis* but seems to rely solely on global regulators for the control of *comK* expression (31–33).

Besides the proximal transcriptional regulation of *comX* by global sensors, competence in *L. lactis* is tightly controlled at the post-transcriptional level through ComX degradation. Indeed, *codY* or *covRS* deletion in three *L. lactis* strains other than DGCC12653 activates *comX* expression at a high level (Fig. S6A) but without unleashing natural transformation. We identified a natural substitution (L125R) in the adaptor protein MecA of strain DGCC12653 that is responsible for its unique behavior regarding spontaneous transformability in laboratory conditions. Moreover, a systematic analysis of more than 600 *L. lactis* genomes confirmed that this MecA variant is unique to this specific strain. This corroborates previous results on strains developing high levels of spontaneous transformation in laboratory conditions that are frequently harboring mutated competence regulators such as CovR in *S. thermophilus* (35) or global repressors in *B. licheniformis* (31). The L125R substitution is located in the linker region between the N- and C-terminal macrodomains of MecA. We previously reported that this region was required for the interaction between MecA and ComX in *S. thermophilus* (62). Thus, we hypothesize that the presence of a charged residue at that position in MecA impairs the interaction with ComX and its subsequent degradation by the MecA-ClpCP machinery. This layer of control through ComX degradation is of major importance for competence development in *L. lactis*, as MecA inactivation in a range of strains boosted spontaneous transformation (Fig. 5C). In *B. subtilis*, the ComS peptide encoded by the *srf* operon activates competence by blocking MecA-ComK interaction, thus avoiding ComK degradation by ClpCP (9, 28, 43, 47, 61). If a similar mechanism is at work in *L. lactis*, it remains to be discovered.

To conclude, our work highlights a different model of competence regulation in *L. lactis* (and possibly *L. cremoris*) than in closely-related streptococci. In this model, the accumulation of the master competence regulator ComX required for natural transformation is under the direct negative control of global nutritional/stress sensors and the Clp degradation machinery. This work paves the way towards a better understanding of the diversity of regulatory networks controlling competence development in Gram-positive bacteria and offers opportunities to use natural transformation as a tool for genome engineering in this lactic acid bacterium of major biotechnological importance.

## MATERIAL AND METHODS

### Bacterial strains, plasmids, and oligonucleotides

Bacterial strains, plasmids, and oligonucleotides used in this study are listed and described in supplemental material (Tables S3-S5).

### Growth conditions

*L. lactis* strains were cultivated in M17 (Difco Laboratories, Detroit, MI) or CDM (adapted from (53)) supplemented with 0.5% (w/v) glucose (M17G and CDMG, respectively), or supplemented with 0.21% (w/v) glucose and 1.1% (w/v) secondary sugar (CDM*) when required, at 30°C without agitation. Glutamate, aspartate, xanthine, adenine, uracil, and guanine were omitted in the competence-optimized CDM (CDM*-DEB). *Escherichia coli* EC1000 was cultivated at 37°C in LB (Lysogeny Broth) medium with agitation. Solid agar plates were prepared by adding 2% (w/v) agar to the medium. When required, 5 μg ml^-1^ of erythromycin, 1 mg ml^-1^ of streptomycin, 0.5 mg ml^-1^ of spectinomycin, and/or 10 μg ml^-1^ of chloramphenicol were added to the medium for *L. lactis*; and 250 μg ml^-1^ of erythromycin, 250 μg ml^-1^ of ampicillin, and 10 μg ml^-1^ of chloramphenicol for *E. coli*.

### DNA techniques and electrotransformation

Electrotransformations of *L. lactis* (30) and *E. coli* (15) were performed as previously described. Chromosomal DNA of *L. lactis* used as template for PCR was extracted as previously described (20). PCRs were performed using the Q5 DNA polymerase (NEB) in a GeneAmp PCR system 2400 (Applied Biosystems).

### Isolation of a *rpsL*\* mutant conferring resistance to streptomycin

Spontaneous streptomycin-resistant IL1403 mutants were isolated on 1 mg ml^-1^ streptomycin-containing plates. After the sequencing of the *rpsL* gene with primers BlD-RpsLUnivUp/BlD-RpsLUnivDown, one spontaneous mutant containing the substitution K56R into the ribosomal protein S12, which was previously shown to confer resistance to streptomycin, was selected (11). A 3.7-kb fragment containing the *rpsL* mutated gene (*strA2* allele, named *rpsL\**) was amplified by PCR with primers BlD-LLcfusARpsL/BlD-LLldacARpsL and cloned into the pGEM-T Easy vector (Promega), yielding plasmid pGEM-*rpsL*\*. This plasmid was used as template to generate the 3.7-kb PCR product with primers BlD-LLcfusARpsL/BlD-LLldacARpsL that was used as donor DNA in natural transformation assays.

### Xylose-induced *comX* expression and natural transformation

The *L. lactis* strains containing plasmid pGhP*_xylT_*-*comX* (P*_xylT_* and *comX* from *L. lactis* IO-1) were grown overnight at 30°C. Cells were washed twice in distilled water, and OD_600_ was adjusted to 0.1 in M17 supplemented with 1% (w/v) xylose and erythromycin. Typically, 2 μg of DNA were added to 100 μl of inoculated transformation medium, and the culture was further incubated during 24 hours at 30°C. Cells were then spread on M17G agar plates supplemented with appropriate antibiotics, and CFUs were counted after 48 hours of incubation at 30°C. The transformation frequency was calculated as the number of antibiotic-resistant CFU ml^-1^ divided by the total number of viable CFU ml^-1^.

### Spontaneous transformation assays

After an overnight preculture in M17G at 30°C of *L. lactis* DGCC12653 (and other strains or mutants), 100 µl of the preculture was diluted in 900 µl of fresh M17G to restart the culture. After 2 hours of growth, cells were washed twice in distilled water and inoculated to a final OD_600_ of 0.1 in competence-activating medium (CDM* or CDM*-DEB). Typically, 2.0 µg of donor DNA were added to 100 µl of transformation medium, and the culture was further incubated for 20 hours at 30°C. Cells were then spread on M17G agar plates supplemented or not with appropriate antibiotics and CFUs were counted after 48 hours of incubation. The transformation frequency was calculated as reported above. The transfer of the mutation conferring streptomycin resistance was confirmed by DNA sequencing of the *rpsL* gene after its amplification by PCR using primers BlD-RpsLUnivUp/BlD-RpsLUnivDown.

### Detection of absorbance and luminescence

Overnight precultures in M17G were washed with sterile water and inoculated in the adequate medium to a final OD_600_ of 0.1. After inoculation in 300 µl-culture volumes and before luminescence analysis, nonyl-aldehyde was diluted 100 times in mineral oil, and 50 µl of this preparation was disposed between the wells of a white 96-well plate with a transparent bottom (Greiner, Alphen a/d Rijn, The Netherlands). Growth (OD_600_) and luciferase activity (Lux) were monitored at 5-minute intervals in a Hidex Sense microplate reader (Hidex, Lemminkäisenkatu, Finland) for a maximum of 24 hours. The luciferase activity is expressed in relative light units (RLU) and the specific luciferase activity in RLU OD_600_ ^-1^.

### Construction of competence reporter plasmids

The P*_comX_-luxAB* reporter plasmid for luminescence assays was constructed as follows. The *comX* promoter was amplified by PCR from chromosomal DNA of *L. lactis* DGCC12653 with primers FT886_PcomX_fw/FT887_PcomX_rec_rv. The plasmid containing the *luxAB* genes pGhP*_comGA[MG]_luxAB* (11) was amplified by PCR using primers FT884_pGhPcomXlux_fw/FT925_PcomXlux_rec_rv. Both PCR fragments were joined by the Gibson assembly method (24) and the resulting plasmid was named pGhP*_comX_*-*luxAB.* The P*_comGA_-luxAB* reporter plasmid for luminescence assays was constructed as follows. The *comGA* promoter was amplified by PCR from chromosomal DNA of *L. lactis* IO-1 using primers LuxcIOF1_XhoI/LuxIOR1. The plasmid pGhP*_comGA[MG]_luxAB* was amplified by PCR using primers LuxcIOR2_XhoI/LuxIOF2. Both PCR fragments were joined by overlap PCR, digested by XhoI, and self-ligated using T4 DNA ligase. The resulting plasmid was named pGhP*_comGA[IO]_*-*luxAB*. The P*_comX_-gfp^sf^* reporter plasmid for fluorescence microscopy was constructed as follows. The *comX* promoter was amplified by PCR from chromosomal DNA of *L. lactis* DGCC12653 using primers FT886_PcomX_fw/FT968_sfGFP_rec. The *gfp^sf^* gene was amplified by PCR from plasmid DNA pDR111_sfGFP(Bs) (44) using primers FT967_sfGFP_fw/FT969_sfGFP_rv_SacII. The plasmid pG^+^host9 was amplified using primers FT970_pGh_SacII/FT925_PcomX_rec_rv. The three PCR fragments were joined by overlapping PCR, digested using SacII, and self-ligated using T4 DNA ligase. The resulting plasmid was named pGhP*_comX_*-*gfp^sf^*.

### Construction of deletion mutants in *L. lactis* strains

The *comX*, *comEC*, *ccpA*, *codY*, *covRS*, and *mecA* genes as well as Rgg and TCS encoding genes were similarly inactivated by the exchange of their coding sequences with either P_32_-*cat* or *spc* resistance cassette (conferring chloramphenicol or spectinomycin resistance, respectively) using double crossing-over events. For this purpose, overlapping PCR products containing the resistance cassette flanked by two recombination arms of ∼1.2 kb (upstream and downstream homologous regions) were generated as previously reported (11). Briefly, upstream, downstream, and resistance-cassette fragments were separately amplified by PCR, purified, mixed in equimolecular amount, and assembled by overlapping PCR by using the most external primers (see Table S5). Typically, 5 µg of the obtained overlapping PCR product were used as donor DNA for xylose-induced natural transformation of the *L. lactis* strains harboring plasmid pGhP*_xylT_-comX*. The correct insertion of the resistance cassette in each targeted locus of the transformants was validated by PCR (see

### Erreur ! Source du renvoi introuvable

To obtain the final mutant strains, the curing of the thermosensitive vector pGhP*_xylT_-comX* was performed by growing the cells 16 hours at 37°C without erythromycin. The cultures were subsequently diluted and plated on M17G agar without erythromycin at 30°C. The resulting colonies were streaked in parallel on M17G plates with and without erythromycin. The absence of plasmid pGhP*_xylT_*-*comX* in erythromycin-sensitive clones was validated by PCR. To generate the double *codY covRS* mutant, *codY* was inactivated by exchange with the *spc* cassette. Then, the high transformability of the Δ*covRS::cat* mutant was used to transfer the Δ*codY::spc* mutation in order to obtain the double mutant.

### Swapping of *mecA* genes

The *mecA* mutants (Δ*mecA::cat*) of strains DGCC12653 and DGCC12671 harboring plasmid pGhP*_xylT_*-*comX* were used for this exchange as they display high transformation rates. The *mecA* genes from strains DGCC12653 and DGCC12671 were amplified by PCR with primers FT1319/FT1320, generating ∼5-kb fragments used as donor DNA. Replacement of the Δ*mecA::cat* deletion by the counterpart *mecA* gene was performed by xylose-induced natural transformation as described above. Five hundred isolated colonies were streaked in parallel on M17G-agar plates containing or not chloramphenicol, searching for antibiotic-sensitive transformants. Then, the swapping of *mecA* genes from sensitive clones was validated by PCR and sequencing. Finally, the curing of the pGhP*_xylT_*-*comX* plasmid was performed as described above.

### Construction of strain DGCC12653 *nisRK* for nisin-induced overexpression

The *nisRK* genes from strain IO-1 were inserted between genes *DGCC12653_06785* and *DGCC12653_06790* using double crossing-over events. For this purpose, an overlapping PCR product containing *nisRK* genes associated with the *spc* cassette in opposite orientation and flanked by two recombination arms of ∼1.2 kb was generated as reported above (see list of primers in Table S5). The obtained overlapping PCR product (5 µg) was used as donor DNA for natural transformation of strain DGCC12653 harboring plasmid pGhP*_xylT_*-*comX*. The correct insertion of *nisRK* genes in the targeted locus of the transformants was validated by PCR. To obtain the final mutant strain, the curing of the thermosensitive vector pGhP*_xylT_*-*comX* was performed as described above.

### Purification of transcriptional regulators

The overexpression plasmids were constructed as follows. The *ccpA*, *codY*, and *covR* genes flanked by an N-terminal 6-His tag encoding sequence were amplified by PCR from strain DGCC12653. For regulator overexpression, pBAD (for *ccpA*) or pNZ8048 (for *codY* and *covR*) was amplified by PCR and joined to the regulator-encoding genes by the Gibson assembly method (see Table S5) (24). The final constructs, pBAD-6his-*ccpA*, pNZ8048-6his-*codY*, and pNZ8048-6his-*covR* were validated by sequencing. The 6His-CcpA (rCcpA) protein was overexpressed in *E. coli* Top10. The 6His-CodY (rCodY) and 6His-CovR (rCovR) were overexpressed in *L. lactis* DGCC12653 *nisRK* (nisin induction with 1 ng ml^-1^). The recombinant proteins were purified using Ni-NTA resin according to the manufacturer’s instructions (ProBond Resin, Novex).

### Electromobility shift assays (EMSA)

The whole promoter region of *comX* (P*_comX_*, intergenic region) from DGCC12653 was amplified by PCR using fluorescent-labeled primers (FT851_Cy3_PcomX_up/FT853_Cy5_PcomX_dw). Shorter versions of P*_comX_* were amplified by PCR using either FT851_Cy3_PcomX_up or FT853_Cy5_PcomX_dw in combination with a non-fluorescent primer located inside P*_comX_* (see list in Table S5). The P*_comX_* fragments harboring the mutated or native version of the CodY-box were amplified by PCR using primer pairs FT1369_PcomX_codY_mute/FT853_Cy5_PcomX_dw and FT1368_PcomX_codY_control/FT853_Cy5_PcomX_dw, respectively. In each assay, the purified proteins were incubated in 2-fold serial dilutions with a constant quantity of labeled probe (2.5 ng µl^-1^). As a negative control, a 150-bp DNA sequence located in the CDS region of the *dnaE* gene was amplified by PCR using primers AK350/AK303 (35). CodY-binding assays were performed in CodY buffer (Tris-HCl 20 mM pH 7.5, NaCl 150 mM, BSA 1 mg ml^-1^, EDTA 1 mM, glycerol 10% [w/v]) at 30°C for 20 minutes. CcpA-binding assays were performed in CodY buffer supplemented with glucose-6-phosphate (25 mg ml^-1^) and poly-dIdC (2 µg ml^-1^) as reported above. CovR-binding assays were performed in Buffer-CovR (NaPO_4_ 50 mM pH 6.5, NaCl 50 mM, MgCl_2_ 1 mM, CaCl_2_ 1 mM, DTT 1 mM, poly-dIdC (2 µg ml^-1^), glycerol 10% [w/v]) at 30°C for 15 minutes. Prior to incubation with the DNA probe, rCovR was incubated in CovR buffer supplemented with 50 mM acetyl phosphate for 30 minutes at 20°C. Samples were migrated in native poly-acrylamide gels in non-denaturing MOPS buffer, and revealed by the Amersham Typhoon device.

### Microscopy

M17G overnight cultures (containing adequate antibiotics) were diluted (1:10) in fresh M17G at 30°C for 2 hours. Cells were then washed in water and inoculated at an OD_600_ of 0.1 in CDM*-DEB until reaching the *comX*-expression peak, determined in parallel with similar clones harboring the pGhP*_comX_*-*luxAB* reporter plasmid. Cells were then centrifuged, resuspended in 50 µl of PBS and observed on agarose pads composed of 1% agarose and PBS buffer (35). Images were obtained using an Axio I inverted microscope (Zeiss) equipped with a Plan-Apochromat objective (100 ×/1.46 oil differential interference contrast [DIC] M27) (Zeiss), a HXP 120 C lighting unit (Zeiss), and a C10600 ORCA-R2 camera (Hamamatsu). GFP fluorescence was detected with filter set 38 HE, displaying bandpass excitation at 470/40 nm and bandpass emission at 525/50 nm (Zeiss). Images were analyzed using ZenPro software (Zeiss) and MicrobeJ (17).

### RNA sequencing of *L. lactis* mutants

RNA extraction and sequencing were performed on the wild type, Δ*codY* mutant, and Δ*covRS* mutant of strain DGCC12653. Each mutant and the wild-type control were grown in their respective efficient competence-activating medium (CDM*-DEB and CDM* for Δ*codY* and Δ*covRS*, respectively) and harvested at their respective diauxic shift. RNA was extracted using the RNeasy plus bacteria kit (Qiagen) according to manufacturer indications. Total RNA was checked for integrity with an RNA Nano chip (Agilent Technologies) and sent for Illumina sequencing (GeneWiz). Raw data were processed on the Galaxy server (use.galaxy.org) using Bowtie2 algorithm to yield BAM files containing the read coordinates and Seqmonk to count the number of reads per coding sequence (CDS). The dataset was exported into an Excel file for further analyses. First, the dataset was standardized to CDS-mapped reads per ∼10 million overall reads. Then, we estimated the ratio of CDS-mapped reads in mutants vs. wild type.

### Data availability

All RNAseq data were deposited in the GEO database under accession number GSEXX The genome of strain DGCC12653 is available under accession number XXXX

## ACKNOWLEDGEMENTS

We warmly thank the technical assistance of Marie-Christine Eloy and Sylvie Derclaye for microscopy experiments. We thank Jan-Willem Veening for the Addgene depository of plasmid pDR111_sfGFP(Bs). This work was supported by the Belgian National Fund for Scientific Research (FNRS, grant PDR T.0110.18) and the Concerted Research Actions (ARC, grant 17/22-084) from Federation Wallonia-Brussels. PH is Research Director at FNRS.

## AUTHOR CONTRIBUTIONS

FT, PhH, CF and PaH conceived and designed the study. FT, MH and FP carried out the laboratory work. FT, J-ML, PhH, CF and PaH analyzed and interpreted the data. FT, J-ML, PhH, CF and PaH wrote or revised the manuscript. All authors read and approved the final manuscript.

