## Supplemental Figures and Tables for "Unveiling the regulatory network controlling natural transformation in lactococci"

**Supplementary Figures:**

**Figure S1. Effects of *comX* overexpression and various glucose-maltose combinations on transformation**

**Figure S2. Effects of CcpA on transformability and *comX* regulation**

**Figure S3. Effects of CodY on *comX* regulation**

**Figure S4. Systematic inactivation of Rgg/TCS**

**Figure S5. Effects of CovRS on *comX* regulation**

**Figure S6. Effects of competence regulators in other *L. lactis* strains**

**Figure S7. $P_{comX}$ activation analyzed by fluorescence microscopy**

**Figure S8. Impact of the double CodY CovRS inactivation on $P_{comX}$ activation**

**Supplementary Tables:**

**Table S1. Up-regulation of competence genes analyzed by RNA sequencing**

**Table S2. Annotation of TCS and Rgg sensors from *L. lactis* DGCC12653**

**Table S3. Strains used and generated in this study**

**Table S4. Plasmids used and generated in this study**

**Table S5. Oligonucleotides used in this study**

**SUPPLEMENTARY FIGURES**

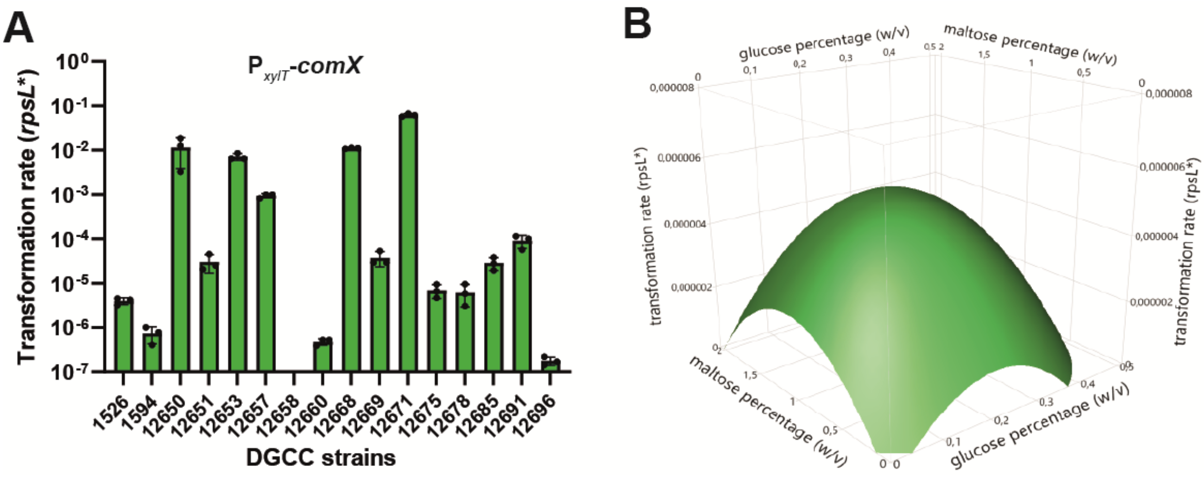


**Figure S1. Effects of *comX* overexpression and various glucose-maltose combinations on transformation.** (A) Transformability upon ComX overexpression. Data show transformation rates of 16 *L. lactis* strains harboring the pGhP$_{xylT}$-*comX* plasmid in M17GX (0.1% glucose-1% xylose). All transformation assays were performed with *rpsL** as donor DNA (20 µg ml$^{-1}$), added at time zero. Cells were spread after ~24 hours of culture. Dots show the values for technical replicates ($n$ = 3) ± standard deviations. (B) Effects of glucose-maltose combinations on spontaneous transformation of strain DGCC12653 WT in CDM. A design of experiment (composite central plan) was setup using the JMP Pro software. Sugar concentrations were varied from 0 to 2% (w/v) in 12 different conditions (glucose %/Maltose %; 0.01/0, 0.05/0.1, 0.3/0.1, 0/0.25, 0.2/0.25, 0.1/0.5, 0.5/0.5, 0.05/1.0, 0.3/1.0, 0/2.0, 0.2/2.0, and 0.4/2.0) containing donor DNA. Compilation of transformation rates was treated with JMP Pro and a response surface was generated.

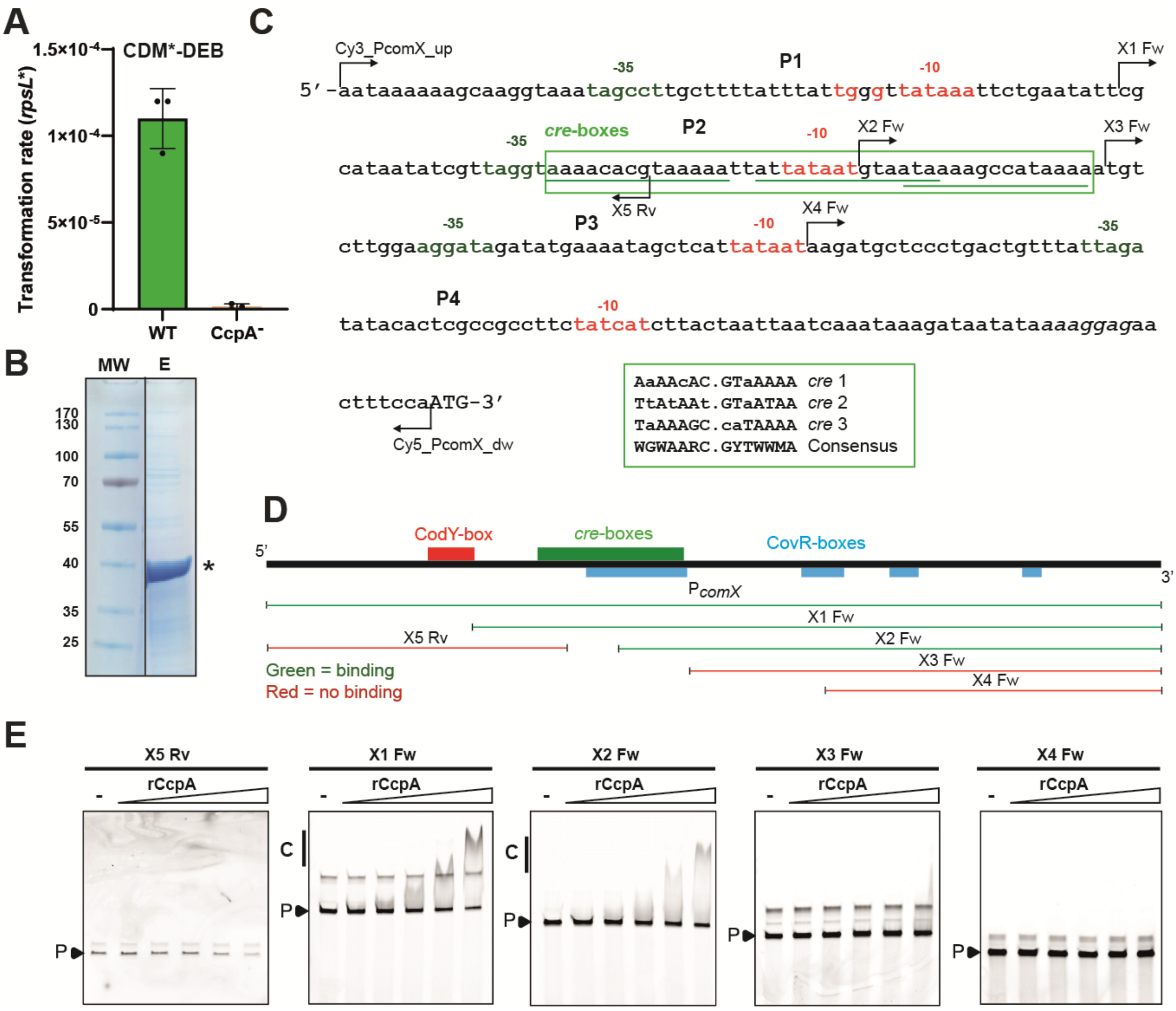


**Figure S2. Effects of CcpA on transformability and *comX* regulation.** (A) Effect of *ccpA* deletion on transformability. Data show transformation rates observed for WT (green) and CcpA$^-$ mutant (light brown) strains in CDM*-DEB after overnight culture with donor DNA. Transformation assays were performed with *rpsL** as donor DNA (20 µg ml$^{-1}$), added at time zero. Cells were spread after ~24 hours of culture. Dots show the values for biological triplicates (CcpA$^-$) or technical triplicates (WT), mean values ± standard deviations. (B) CcpA purification. SDS-PAGE of the elution step [E] of 6His-CcpA (rCcpA) purified from *E. coli*. MW, molecular weight (kDa). The star indicates the enriched rCcpA. (C) Mapping of fluorescent (Cy3_PcomX_up, Cy5_PcomX_dw) and non-fluorescent (X5 Rv, X1 Fw, X3 Fw, X2 Fw, X3 Fw, and X4 Fw) primers designed in $P_{comX}$ (complete intergenic region). The region containing three *cre*-boxes (underlined) is surrounded in green. The alignment of the three *cre*-boxes with the consensus is shown with conserved positions in capitals. (D) Mapping of the different probes used for EMSAs. Green and red lines indicate the presence or absence of a band shift. (E) EMSAs performed with a gradient of purified CcpA (rCcpA) on the different probes shown in panel D. Lanes without rCcpA are indicated by a minus sign. C and P indicate the CcpA-DNA complex(es) and the unbound probe, respectively.

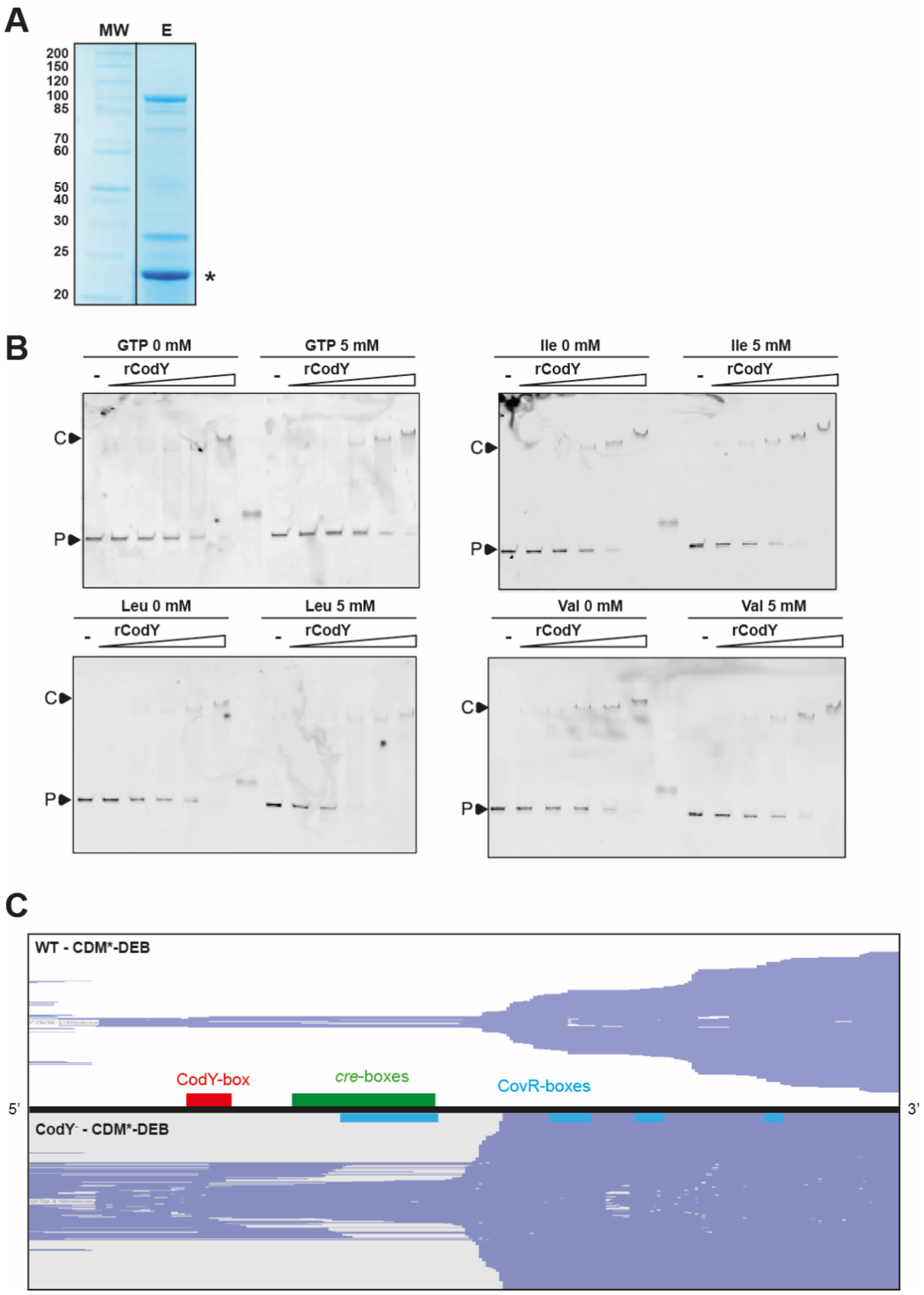


**Figure S3. Effects of CodY on *comX* regulation.** (A) CodY purification. SDS-PAGE of the elution step [E] of the 6His-CodY (rCodY) purified from *L. lactis*. MW, molecular weight (kDa). The star indicates the enriched rCodY. (B) EMSAs performed with a gradient of purified CodY (rCodY) on $P_{comX}$ in absence (left side of each panel) or presence (right side of each panel) of 5 mM of GTP (top left panel), Ile (top right panel), Leu (lower left panel) or Val (lower right panel). Lanes without rCodY are indicated by a minus sign. C and P indicate rCodY-$P_{comX}$ complex and unbound probe, respectively. (C) Mapping of RNAseq reads on $P_{comX}$, from WT (top panel) and $CodY^{-}$ (lower panel) strains. Cells were harvested at the diauxic shift in CDM*-DEB. The $P_{comX}$ (complete intergenic region) is illustrated as a black line, while CodY-, *cre*- and CovR-boxes are localized and displayed in red, green and blue, respectively.

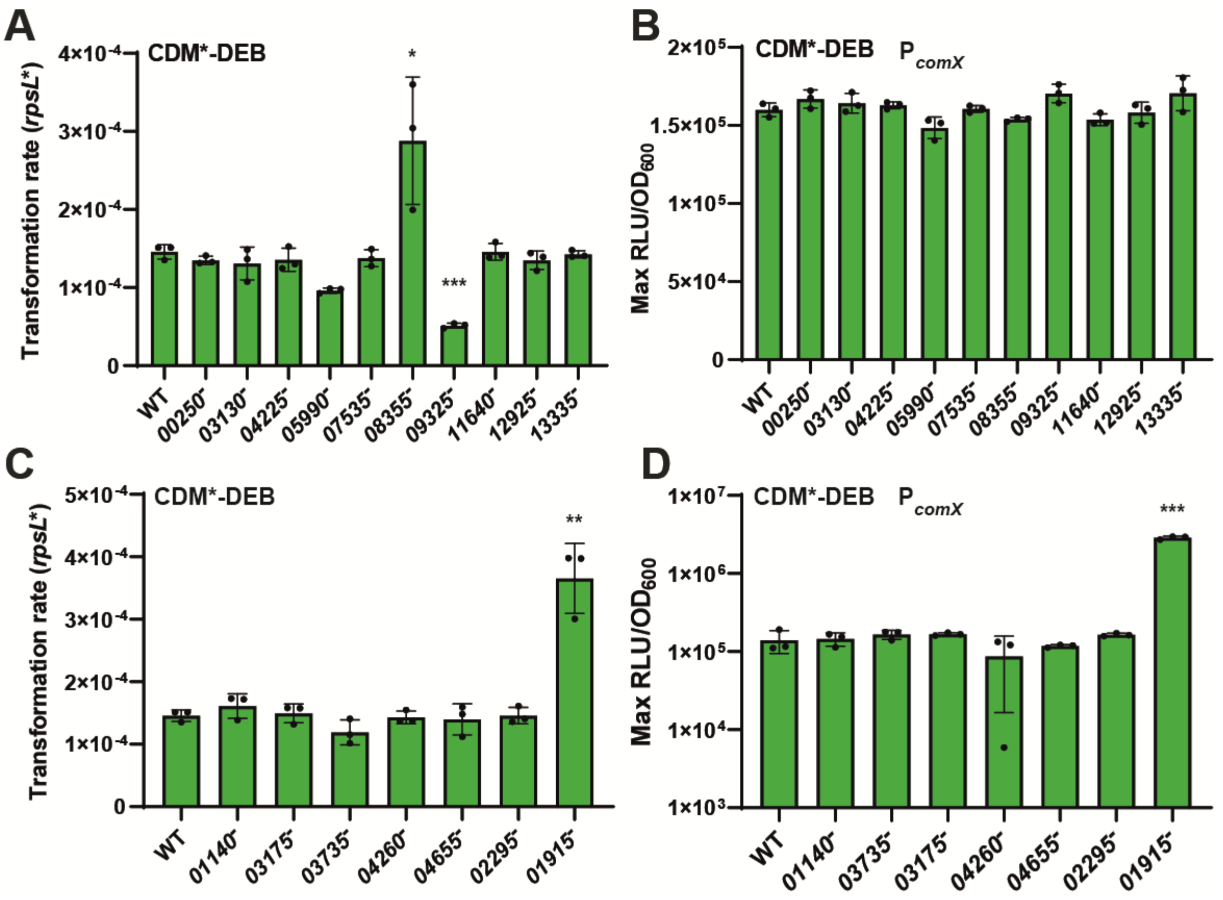


**Figure S4. Systematic inactivation of Rgg/TCS.** (A and B) Effects on transformability (A) and $P_{comX}$ activity (B) of the independent inactivation of 10 Rgg from DGCC12653. (C and D) Effects on transformability (C) and $P_{comX}$ activity (D) of the independent inactivation of 7 TCS from DGCC12653 (labels indicate the first gene of the TCS). Data on transformability (panels A and C) show transformation rates observed for WT and mutant strains in CDM*-DEB after overnight culture with donor DNA. Transformation assays were performed with *rpsL** as donor DNA (20 µg $ml^{-1}$), added at time zero. Cells were spread after ~24 hours of culture. Data on $P_{comX}$ activity (panels B and D) show maximum specific luciferase activity (Max RLU/$OD_{600}$) observed at the diauxic shift in CDM*-DEB. Dots (panels A to D) show the values for biological triplicates ($Rgg^-$ and $TCS^-$ strains) or technical triplicates (WT), mean values ± standard deviations. Statistical *t* test was performed for each mutant strain in comparison to the WT ($n$=3; *, $P < 0.05$; **, $P < 0.01$; ***, $P < 0.001$).

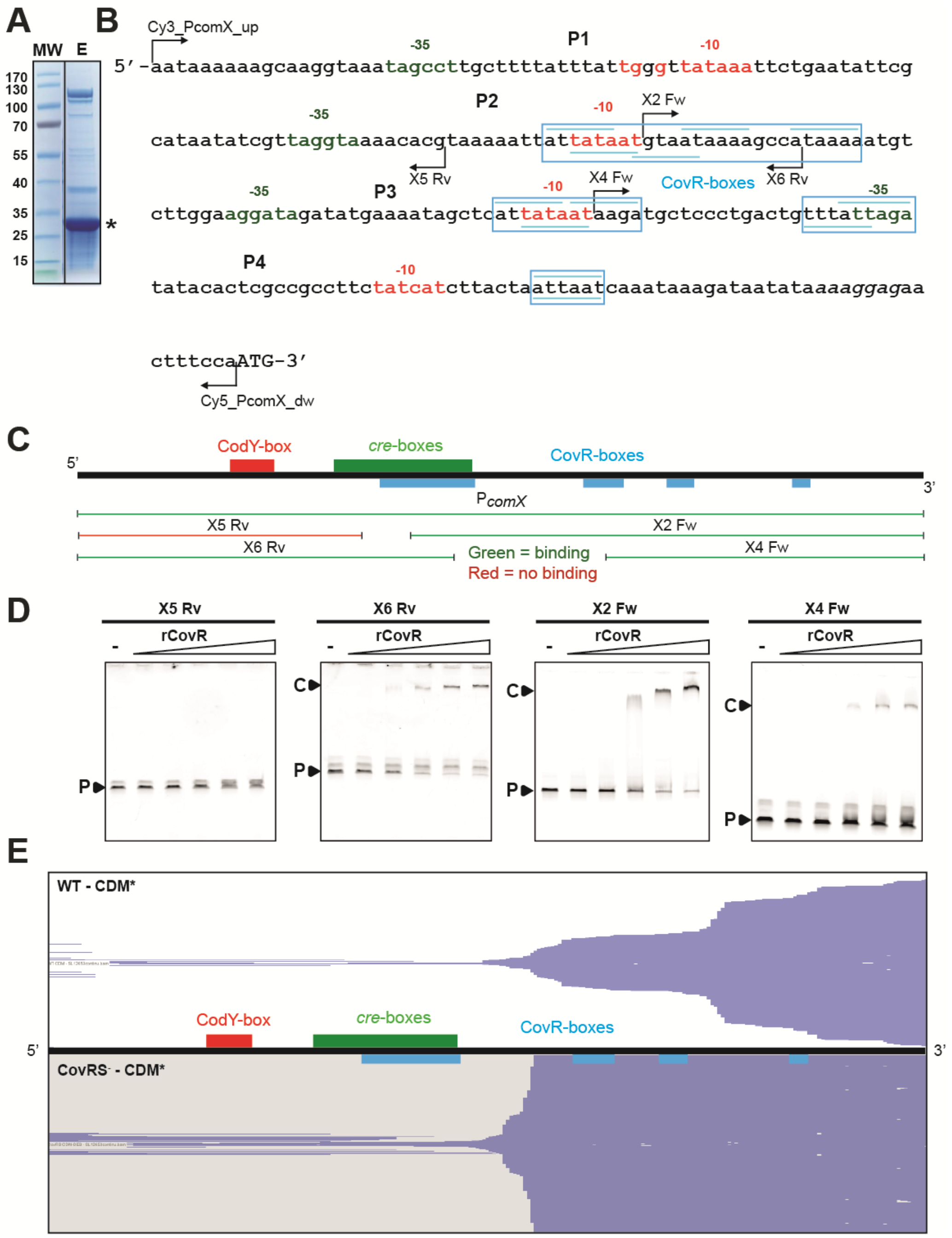


**Figure S5. Effects of CovRS on *comX* regulation.** (A) CovR purification. SDS-PAGE of the elution step [E] of the 6His-CovR (rCovR) purified from *L. lactis*. MW, molecular weight (kDa). The star indicates the enriched rCovR. (B) Mapping of fluorescent (Cy3_PcomX_up, Cy5_PcomX_dw) and non-fluorescent (X5 Rv, X6 Rv, X2 Fw, X3 Fw, and X4 Fw) primers designed in $P_{comX}$ (complete intergenic region). The regions containing CovR-boxes (underlined) are surrounded in blue. (C) Mapping of the different probes used for EMSAs. Green and red lines indicate the presence or absence of a band shift. (D) EMSAs performed with a gradient of purified CovR (rCovR) on the different probes shown in panel C. Lanes without rCovR are indicated by a minus sign. C and P indicate the rCovR-DNA complex(es) and the unbound probe, respectively. (E) Mapping of RNAseq reads on $P_{comX}$, from WT (top panel) and CovRS$^-$ (lower panel) strains. Cells were harvested at the diauxic shift in CDM*. The $P_{comX}$ (complete intergenic region) is illustrated as a black line, while CodY-, *cre*- and CovR-boxes are localized and displayed in red, green, and blue, respectively.

**A**

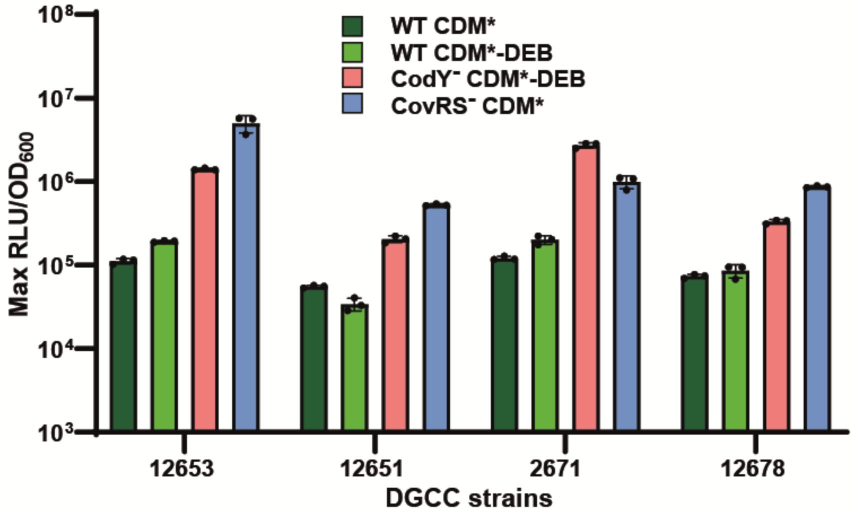


**B**

```
12653_04910      MKYEDINENTIKITLSFDDLTDYDIKLSDFFGNQEVIEQFFYELVDELGLENRFGNVGML   60
12651_03500      MKYEDINENTIKITLSFDDLTDYDIKLSDFFGNQEVIEQFFYELVDELGLENRFGNVGML   60
12671_01635      MKYEDINENTIKITLSFDDLTDYDIKLSDFFGNQEVIEQFFYELVDELGLENRFGNVGML   60
12678_05760      MKYEDINENTIKITLSFDDLTDYDIKLSDFFGNQEVIEQFFYELVDELGLENRFGNVGML   60
                 ************************************************************

12653_04910      TFQIQPFPQGVHMIVHEEAMLGEGGEIPDDPEEFEELMTGFYNKLNEIGADMARERGITD   120
12651_03500      TFQIQPFPQGVHMIVHEEAMLGEGGEIPDDPEEFEELMTGFYNKLNEIGADMARERGITD   120
12671_01635      TFQIQPFPQGVHMIVHEEAMLGEGGEIPDDPEEFEELMTGFYNKLNEIGADMARERGITD   120
12678_05760      TFQIQPFPQGVHMIVHEEAMLGEGGEIPDDPEEFEELMTGFYNKLNEIGADMARERGITD   120
                 ************************************************************

12653_04910      FKPGRGLPGTKKDEAEQEPDFIYYSIRYEDIMSVLTGIKNVKFADEESEFYRYDGNFYLV   180
12651_03500      FKPGLGLPGTKKDEAEQEPDFIYYSIRYEDIMSVLTGIKNVKFADEESEFYRYDGNFYLV   180
12671_01635      FKPGLGLPGTKKDEAEQEPDFIYYSIRYEDIMSVLTGIKNVKFADEESEFYRYDGNFYLV   180
12678_05760      FKPGLGLPGTKKDEAEQEPDFIYYSIRYEDIMSVLTGIKNVKFADEESEFYRYDGNFYLV   180
                 **** *******************************************************

12653_04910      VLDNQKEKGKMHVESTRSRMMEYGEATKMSREFLQEYGECLIATRALDVLRKI   233
12651_03500      VLDNQKEKGKMHVESTRSRMMEYGEATKMSREFLQEYGECLIATRALDVLRKI   233
12671_01635      VLDNQKEKGKMHVESTRSRMMEYGEATKMSREFLQEYGECLIATRALDVLRKI   233
12678_05760      VLDNQKEKGKMHVESTRSRMMEYGEATKMSREFLQEYGECLIATRALDVLRKI   233
                 *****************************************************
```

**Figure S6. Effects of competence regulators in other *L. lactis* strains.** (A) Effects of CodY and CovRS inactivation on $P_{comX}$ activity in strains DGCC12653, DGCC12651, DGCC12671 and DGCC12678. The effects of these deletions ($CodY^-$ in CDM*-DEB, light red bars and $CovRS^-$ in CDM*, blue bars) were compared to their respective WT in the same culture conditions (WT in CDM*, dark green bars and CDM*-DEB, light green bars). Data show maximum specific luciferase activity (Max $RLU/OD_{600}$) observed at the diauxic shift either in CDM*-DEB or CDM*. Dots show the values for biological triplicates ($CodY^-$ and $CovR^-$ strains) or technical triplicates (WT), mean values ± standard deviations. (B) Sequence alignment of MecA proteins from strains DGCC12653, DGCC12651, DGCC12671 and DGCC12678. The R125 residue is in red and underlined. The alignment was performed using Clustal Omega.

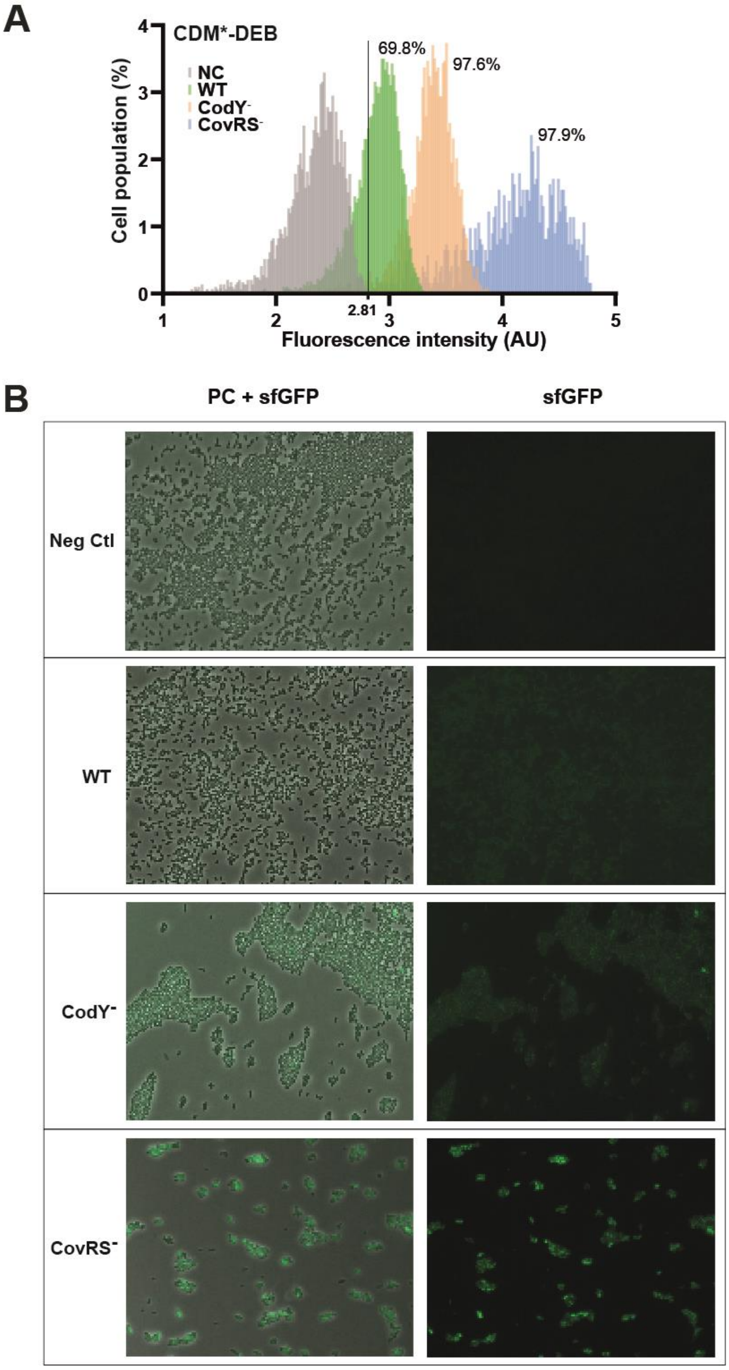


**Figure S7. $P_{comX}$ activation analyzed by fluorescence microscopy.** (A) Density plot of single-cell fluorescence intensity (arbitrary unit [AU]) for WT, CodY$^-$, CovRS$^-$ mutant harboring the reporter fusion $P_{comX}$-*gfp*$^{sf}$ (green, orange and blue, respectively) and the negative control (NC) harboring the empty plasmid (grey). Cells were cultured in CDM*-DEB, harvested at the diauxic shift, and analyzed by epifluorescence microscopy. The fluorescence of more than 1,200 individual cells was examined in each experiment. A minimum threshold of significant fluorescence (2.81 AU) was determined as the maximum signal reached by the negative control. Percentages in the top of the plot indicate the percentage of the cell population displaying a fluorescence signal higher than 2.81 for each strain. (B) Representative images of experiments depicted in panel A for two channels (left, merge phase contrast [PC] and sfGFP signal; and right, sfGFP signal).

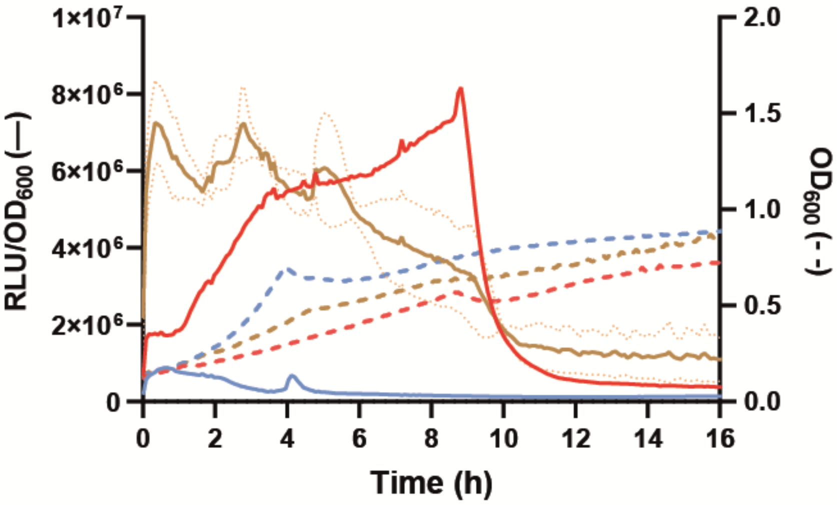


**Figure S8. Impact of the double CodY CovRS inactivation on $P_{comX}$ activation.** Growth (dotted lines – $OD_{600}$) and kinetics of $P_{comX}$ specific luciferase activity (continuous lines – RLU/$OD_{600}$) monitored over time for CodY$^-$ (blue), CovRS$^-$ (red) and CodY$^-$ CovRS$^-$ (brown) mutant strains. Continuous and dotted dark lines are the mean values of biological triplicates, and light lines are standard deviations.

### SUPPLEMENTARY TABLES

**Table S1. Up-regulation of competence genes analyzed by RNA sequencing**

| | Locus tag | Strand | Gene | Description | $\frac{\Delta codY}{WT}$ | $\frac{\Delta covRS}{WT}$ | $\frac{PxylTcomX}{WT}$ |
|---|---|---|---|---|---|---|---|
| | DGCC12653_13390 | + | *comX* | competence factor | 8.19 | 5.97 | 207.78 |
| Pore assembly | DGCC12653_09995 | + | *comGA* | Late competence protein ComGA, access of DNA to ComEA | 23.46 | 3305.56 | 23445.92 |
| | DGCC12653_10000 | + | *comGB* | Late competence protein ComGB, access of DNA to ComEA | 23.84 | 1327.44 | 53635.43 |
| | DGCC12653_10005 | + | *comGC* | Late competence protein ComGC, access of DNA to ComEA | 29.94 | 622.61 | 6077.95 |
| | DGCC12653_10010 | + | *comGD* | Late competence protein ComGD, access of DNA to ComEA | 21.71 | 146.84 | 3967.67 |
| | DGCC12653_10015 | + | *comGE* | Late competence protein ComGE | 16.19 | 251.91 | 8578.22 |
| | DGCC12653_10020 | + | *comGF* | Late competence protein ComGF, access of DNA to ComEA | 29.79 | 542.33 | 12979.23 |
| | DGCC12653_10025 | + | *comGG* | Late competence protein ComGG | 25.57 | 18.93 | 619.13 |
| | DGCC12653_09640 | + | *comC* | Late competence protein ComC, processing protease | 18.73 | 209.62 | 573.93 |
| DNA uptake | DGCC12653_04795 | - | *comEA* | Late competence protein ComEA, DNA receptor | 17.86 | 336.76 | 1281.01 |
| | DGCC12653_04790 | - | *comEC* | Late competence protein ComEC, DNA transport | 19.96 | 183.82 | 1824.13 |
| | DGCC12653_02995 | + | *comFA* | Late competence protein ComFA, DNA transporter ATPase | 29.07 | 354.24 | 1934.04 |
| | DGCC12653_03000 | + | *comFC* | Late competence protein ComFC, phosphorybosyltransferase domain | 28.64 | 2362.51 | 3061.24 |
| Recombination | DGCC12653_06395 | + | *ssbA* | ssDNA binding protein | 17.49 | 178.41 | 1164.19 |
| | DGCC12653_02130 | - | *dprA* | DNA recombination-mediator protein A | 26.86 | 683.19 | 20923.49 |
| | DGCC12653_04550 | - | *coiA* | Competence protein CoiA | 16.32 | 21.59 | 41.42 |
| | DGCC12653_06225 | + | *recA* | Recombinase A protein | 2.62 | 3.16 | 13.41 |

**Table S2. Annotation of TCS and Rgg sensors from *L. lactis* DGCC12653**

| | *L. lactis* DGCC12653 | | | | *L. lactis* KF147 |
|---|---|---|---|---|---|
| | **Locus tag** | **Initial gene name[§]** | **Revised gene name** | **Product** | **Locus tag** |
| Two-Component Systems | DGCC12653_01915 | *llrA* | *covR* | RR CovR | LLKF_1744 |
| | DGCC12653_01910 | | *covS* | KH CovS | LLKF_1743 |
| | DGCC12653_01140 | *llrB* | *sphR* | RR SphR | LLKF_1540 |
| | DGCC12653_01145 | | *sphS* | HK SphS | LLKF_1541 |
| | DGCC12653_06420 | *llrC* | *walR/vicR* | RR VicR | LLKF_0449 |
| | DGCC12653_06415 | | *walH/vicK* | HK VicK | LLKF_0448 |
| | DGCC12653_03735 | *llrD* | *vraR* | RR VraR | LLKF_0918 |
| | DGCC12653_03740 | | *vraS* | HK VraS | LLKF_0917 |
| | DGCC12653_03175 | *llrE* | *llrE* | RR LlrE | LLKF_1024 |
| | DGCC12653_03165 | | *kinE* | HK KinE | LLKF_1026 |
| | DGCC12653_04260 | *llrF* | *ciaR* | RR CiaR | LLKF_1834 |
| | DGCC12653_04255 | | *ciaH* | HK CiaH | LLKF_1833 |
| | DGCC12653_04655 | *llrG* | *llrG* | RR LlrG | LLKF_1918 |
| | DGCC12653_04650 | | *kinG* | HK KinG | LLKF_1917 |
| | DGCC12653_02295 | *kdpED* | *kdpE* | RR KdpE | LLKF_1239 |
| | DGCC12653_02300 | | *kdpD* | HK KdpD | LLKF_1238 |
| Rgg sensors | DGCC12653_00250 | | *gadR* | GadR | LLKF_1358 |
| | DGCC12653_03130 | | *ykhI* | YkhI | LLKF_1092 |
| | DGCC12653_04225* | | *yrbI* | YrbI | LLKF_1827 |
| | DGCC12653_05990 | | *ywdE_bis* | YwdE_bis | LLKF_0365 |
| | DGCC12653_07535 | | *yhgC* | YhgC | LLKF_0776 |
| | DGCC12653_08355 | | *yrbI_bis* | YrbI_bis | LLKF_0051 |
| | DGCC12653_09325 | | *yueB* | YueB | NH[#] |
| | DGCC12653_11640 | | *ywiI* | YwiI | LLKF_2477 |
| | DGCC12653_12925 | | *(kw2_0044)* | Kw2_0044 | LLKF_0034† |
| | DGCC12653_13335 | | *ywdE* | YwdE | LLKF_2403 |

§ gene name according to *Lactococcus cremoris* MG1363 (5)
* inactive, misplaced start codon
† truncated, only N-terminal part
\# NH, No Homolog

**Table S3. Strains used and generated in this study**

| Species/Strain | Characteristics | Source |
|---|---|---|
| *E. coli* | | |
| Top 10 | $F^-$ *mcrA* Δ(*mrr hsdRMS-mcrBC*) φ80*lac*ZΔM15 Δ*lacX74 recA1 deoR ara*D139 Δ(*ara-leu*)7697 *galU galK rpsL endA1 nupG* | Invitrogen |
| *L. lactis* | | |
| 1526 | Wild-type dairy isolate | IFF/Danisco collection |
| DGCC1594 | Wild-type dairy isolate | IFF/Danisco collection |
| DGCC12650 | Wild-type plant isolate | IFF/Danisco collection |
| DGCC12651 | Wild-type plant isolate | IFF/Danisco collection |
| DGCC12653 | Wild-type plant isolate | IFF/Danisco collection |
| DGCC12657 | Wild-type plant isolate | IFF/Danisco collection |
| DGCC12658 | Wild-type plant isolate | IFF/Danisco collection |
| DGCC12660 | Wild-type plant isolate | IFF/Danisco collection |
| DGCC12662 | Wild-type plant isolate | IFF/Danisco collection |
| DGCC12668 | Wild-type plant isolate | IFF/Danisco collection |
| DGCC12669 | Wild-type plant isolate | IFF/Danisco collection |
| DGCC12671 | Wild-type plant isolate | IFF/Danisco collection |
| DGCC12675 | Wild-type plant isolate | IFF/Danisco collection |
| DGCC12678 | Wild-type plant isolate | IFF/Danisco collection |
| DGCC12685 | Wild-type plant isolate | IFF/Danisco collection |
| DGCC12686 | Wild-type plant isolate | IFF/Danisco collection |
| DGCC12691 | Wild-type plant isolate | IFF/Danisco collection |
| DGCC12696 | Wild-type plant isolate | IFF/Danisco collection |
| FRT101 | DGCC12653 *codY*::$P_{32}$-*cat* | This study |
| FRT102 | DGCC12653 *covRS*::$P_{32}$-*cat* (*01910-01915*) | This study |
| FRT103 | DGCC12653 *mecA*::$P_{32}$-*cat* | This study |
| FRT104 | DGCC12653 *ccpA*::$P_{32}$-*cat* | This study |
| FRT105 | DGCC12653 *covRS*::$P_{32}$-*cat codY*::*spec* | This study |
| FRT106 | DGCC12653 *comX*::$P_{32}$-*cat* | This study |
| FRT107 | DGCC12653 *comEC*::$P_{32}$-*cat* | This study |
| FRT108 | DGCC12653 *mecA* (L-125) | This study |
| FRT109 | DGCC12671 *mecA* (R-125) | This study |
| FRT201 | DGCC1594 *mecA*::$P_{32}$-*cat* | This study |
| FRT202 | DGCC12650 *mecA*::$P_{32}$-*cat* | This study |
| FRT203 | DGCC12651 *mecA*::$P_{32}$-*cat* | This study |
| FRT204 | DGCC12657 *mecA*::$P_{32}$-*cat* | This study |
| FRT205 | DGCC12660 *mecA*::$P_{32}$-*cat* | This study |
| FRT206 | DGCC12669 *mecA*::$P_{32}$-*cat* | This study |
| FRT207 | DGCC12671 *mecA*::$P_{32}$-*cat* | This study |
| FRT208 | DGCC12675 *mecA*::$P_{32}$-*cat* | This study |
| FRT209 | DGCC12678 *mecA*::$P_{32}$-*cat* | This study |
| FRT210 | DGCC12685 *mecA*::$P_{32}$-*cat* | This study |
| FRT211 | DGCC12691 *mecA*::$P_{32}$-*cat* | This study |
| FRT211 | DGCC12651 *codY*::$P_{32}$-*cat* | This study |
| FRT212 | DGCC12651 *covRS*::$P_{32}$-*cat* | This study |
| FRT213 | DGCC12671 *codY*::$P_{32}$-*cat* | This study |
| FRT214 | DGCC12671 *covRS*::$P_{32}$-*cat* | This study |
| FRT215 | DGCC12678 *codY*::$P_{32}$-*cat* | This study |
| FRT216 | DGCC12678 *covRS*::$P_{32}$-*cat* | This study |
| FRT217 | DGCC12653 *ecto::nisRK-spc* | This study |
| FRT218 | DGCC12653 *00250*::*spc* | This study |
| FRT219 | DGCC12653 *03130*::*spc* | This study |
| FRT220 | DGCC12653 *04225*::*spc* | This study |
| FRT221 | DGCC12653 *05990*::*spc* | This study |
| FRT222 | DGCC12653 *07535*::*spc* | This study |
| FRT223 | DGCC12653 *08355*::*spc* | This study |
| FRT224 | DGCC12653 *09325*::*spc* | This study |
| FRT225 | DGCC12653 *11640*::*spc* | This study |
| FRT226 | DGCC12653 *12925*::*spc* | This study |
| FRT227 | DGCC12653 *13335*::*spc* | This study |
| FRT228 | DGCC12653 *01140-01145*::$P_{32}$-*cat* | This study |
| FRT229 | DGCC12653 *03165-03175*::$P_{32}$-*cat* | This study |
| FRT230 | DGCC12653 *03735-03740*::$P_{32}$-*cat* | This study |
| FRT231 | DGCC12653 *04255-04260*::$P_{32}$-*cat* | This study |
| FRT232 | DGCC12653 *04650-04655*::$P_{32}$-*cat* | This study |
| FRT233 | DGCC12653 *02295-02300*::$P_{32}$-*cat* | This study |

### Table S4. Plasmids used and generated in this study

| Plasmid | Characteristics | Source or reference |
|---|---|---|
| pBAD/His | $Ap^r$; expression vector | Invitrogen |
| pBAD_6his-*ccpA* | $Ap^r$; pBAD derivative carrying 6his-*ccpA* under the control of the arabinose-inducible promoter $P_{araB}$ | This study |
| $pG^+$host9 | $Em^r$ Ts | (4) |
| pNZ5319 | $Em^r$ $Cm^r$; pACYC184 derivative containing the $P_{32}$-*cat* cassette surrounded by *lox* sites | (3) |
| pJUD-*spc* | $Em^r$ $Spc^r$; $pG^+$host9 derivative carrying the *spc* cassette | L. Fontaine (laboratory collection) |
| pJIM4900 | $Em^r$ Ts; $pG^+$host9 derivative containing the *luxAB* genes of *Photorhabdus luminescens* | E. Guédon (laboratory collection) |
| pGh$P_{xylT}$-*comX* | $Em^r$ Ts; $pG^+$host9 derivative carrying *comX* under the control of the inducible promoter $P_{xylT}$, both cloned from *L. lactis* IO-1 | (1) |
| pGh$P_{comX}$-*luxAB* | $Em^r$ Ts; $pG^+$host9 derivative carrying *luxAB* genes under the control of the early promoter $P_{comX}$ of *L. lactis* DGCC12653 | This study |
| pGh$P_{comGA}$-*luxAB* | $Em^r$ Ts; $pG^+$host9 derivative carrying *luxAB* genes under the control of the late promoter $P_{comGA}$ of *L. lactis* IO-1 | This study |
| pGh$P_{comX}$-$gfp^{sf}$ | $Em^r$ Ts; $pG^+$host9 derivative carrying $gfp^{sf}$ genes under the control of the early promoter $P_{comX}$ of *L. lactis* DGCC12653 | This study |
| pDR111_$gfp^{sf}$(*Bs*) | *bla amyE'* $P_{hyperspank}$-*sfgfp*(*Bs*) *spc lacI 'amyE* | (6) |
| pGEM-*rpsL** | $Ap^r$; pGEM-T Easy derivative carrying the *rpsL** gene from *L. lactis* IL1403 | This study |
| pNZ8048 | $Cm^r$; Translational fusion vector carrying $P_{nisA}$ + terminator | (2) |
| pNZ8048_6his-*codY* | $Cm^r$; pNZ8048 derivative carrying 6his-*codY* under the control of the nisin-inducible promoter $P_{nisA}$ | This study |
| pNZ8048_6his-*covR* | $Cm^r$; pNZ8048 derivative carrying 6his-*covR* under the control of the nisin-inducible promoter $P_{nisA}$ | This study |

$Ap^r$, $Em^r$, $Cm^r$, and $Spc^r$: ampicillin, erythromycin, chloramphenicol, and spectinomycin resistance, respectively
Ts: thermosensitive

### Table S5. Oligonucleotides used in this study

| Target fragment | Primer name | Primer sequence (5'-3') | Template DNA | PCR fragment size |
|---|---|---|---|---|
| *codY*::P32-*cat* | | | | |
| Up recombination arm | FT767_codY_locus_fw | CTTTTGGACACAGGGGATGAGG | DGCC12653 chromosome | 1148 bp |
| | FT768_codY_rec_rv | GCCCTTATGGGATTTATCTTCCTTATCAGTCATACTG TTATATGGCAACTCTTGT | | |
| Resistance cassette | 252-Uplox66 | TAAGGAAGATAAATCCCATA | pNZ5319 | 1402 bp |
| | 253-DNlox71 | TTCACGTTACTAAAGGGAATGTA | | |
| Down recombination arm | FT769_codY_rec_fw | TCTACATTCCCTTTAGTAACGTGAAACTGGTTTATTT GATAAACTTGCAGGACGT | DGCC12653 chromosome | 1225 bp |
| | FT770_codY_locus_rv | GTTATTGTATGACATTAAGCATC | | |
| Insertion validation | 252-Uplox66 | TAAGGAAGATAAATCCCATA | clones | 2800 bp |
| | FT771_codY_diagR | TCCGTCAGTCACAACGTCTGG | | |
| *covRS*::P32-*cat* (*01910-01915*::P32-*cat*) | | | | |
| Up recombination arm | FT732_01915_locus_fw | CAACTTTGGTCTGGCAAACAGTCC | DGCC12653 chromosome | 1223 bp |
| | FT733_019115_rec_rv | GCCCTTATGGGATTTATCTTCCTTAGTCGCATAGCCT TCATGTTCTAATTCTAAT | | |
| Resistance cassette | 252-Uplox66 | TAAGGAAGATAAATCCCATA | pNZ5319 | 1402 bp |
| | 253-DNlox71 | TTCACGTTACTAAAGGGAATGTA | | |
| Down recombination arm | FT734_01915_rec_fw | TCTACATTCCCTTTAGTAACGTGAATTTAGCTCGATT GGCAGAAAATTATCAAGG | DGCC12653 chromosome | 1272 bp |
| | FT735_01915_locus_rv | ATCCCGCCAATACCGTGACAAC | | |
| Insertion validation | 252-Uplox66 | TAAGGAAGATAAATCCCATA | clones | 2805 bp |
| | FT736_01915_diagR | ATGACCCAACTCACAATAGC | | |
| *mecA*::P32-*cat* | | | | |
| Up recombination arm | FT681_mecA_locus_fw | TTTAATCCTTGCAAAGAAGCAACCGAGA | DGCC12653 chromosome | 1104 bp |
| | FT 679_mecA_rec_fw | CGCCCTTATGGGATTTATCTTCCTTACGTTATGAAGA TATTATGTCTGTCTTAAC | | |
| Resistance cassette | 252-Uplox66 | TAAGGAAGATAAATCCCATA | pNZ5319 | 1402 bp |
| | 253-DNlox71 | TTCACGTTACTAAAGGGAATGTA | | |
| Down recombination arm | FT680_mecA_rec_fw | TCTACATTCCCTTTAGTAACGTGAATATCACTTTACC AGTCTTATTAGTATTATA | DGCC12653 chromosome | 1100 bp |
| | FT682_mecA_locus_rv | CTTTGAGAAACATTCTACCAGCCGTC | | |
| Insertion validation | 252-Uplox66 | TAAGGAAGATAAATCCCATA | clones | 2713 bp |
| | FT683_mecA_diagR | TCTCATATTATAAAAGCCAGTCATTAG | | |
| *ccpA*::P32-*cat* | | | | |
| Up recombination arm | FT720_ccpA_locus_fw | CGTACCCCTATTTAGAATACGC | DGCC12653 chromosome | 1226 bp |
| | FT719_ccpA_DGCC_rec_rv | GCCCTTATGGGATTTATCTTCCTTAAGAAGATGTTG AACAAAATCAATTAGTTTTGG | | |
| Resistance cassette | 252-Uplox66 | TAAGGAAGATAAATCCCATA | pNZ5319 | 1402 bp |
| | 253-DNlox71 | TTCACGTTACTAAAGGGAATGTA | | |
| Down recombination arm | FT634_CcpA_kw2_rec_fw | CTACATTCCCTTTAGTAACGTGAACATAGTATTTATG AAAACCATTTTCATTTAC | DGCC12653 chromosome | 1276 bp |
| | FT635_CcpA_kw2_locus_r v | GGTCAGTATGAGTGAAACTTTCACAAC | | |
| Insertion validation | 252-Uplox66 | TAAGGAAGATAAATCCCATA | clones | 2941 bp |
| | FT721_ccpA_diagR | ATGCACCAGATGGTCGGTA | | |
| *codY*::*spc* | | | | |
| Up recombination arm | FT767_codY_locus_fw | CTTTTGGACACAGGGGATGAGG | DGCC12653 chromosome | 1148 bp |
| | FT768_codY_rec_rv | GCCCTTATGGGATTTATCTTCCTTATCAGTCATACTG TTATATGGCAACTCTTGT | | |
| Resistance cassette | 252-Uplox66 | TAAGGAAGATAAATCCCATA | pJUD-*spc* | 1034 bp |
| | 253-DNlox71 | TTCACGTTACTAAAGGGAATGTA | | |

| | | | | |
|---|---|---|---|---|
| Down recombination arm | FT769_codY_rec_fw | TCTACATTCCCTTTAGTAACGTGAAACTGGTTTATTTGATAAACTTGCAGGACGT | DGCC12653 chromosome | 1225 bp |
| | FT770_codY_locus_rv | GTTATTGTATGACATTAAGCATC | | |
| Insertion validation | 252-Uplox66 | TAAGGAAGATAAATCCCATA | clones | 2432 bp |
| | FT771_codY_diagR | TCCGTCAGTCACAACGTCTGG | | |
| *ecto*::*nisRK-spc* | | | | |
| Up recombination arm | FT929_ecto_1500_fw | CTTGCTTCTTGATAAAGGAAAGTTGCA | DGCC12653 chromosome | 1527 bp |
| | FT1125_nisRK_ecto_rec_rv | TGATATTAAATCTGGAACAGTCTGTGGGCCGCCCAATATGACAAGAGCGACTAT | | |
| *nisRK* genes | FT1055_nisRK_aval | CCACAGACTGTTCCAGATTTAATATCA | IO-1 chromosome | 2665 bp |
| | FT1057_nisRK_spec_rec | ATCCTTATGGGATTTATCTTCCTTAAAGCTTTTAGCTTAGATACAGATAAAG | | |
| Resistance cassette | 252-Uplox66 | TAAGGAAGATAAATCCCATA | pJUD-*spc* | 1034 bp |
| | 253-DNlox71 | TTCACGTTACTAAAGGGAATGTA | | |
| Down recombination arm | FT711_ectopic_rec_fw | ATCTACATTCCCTTTAGTAACGTGAAATACCAGTTTGACTTTACCAAAGTATAGTG | DGCC12653 chromosome | 1523 bp |
| | FT934_ecto_1500_rv | AAATTTGTAAGCAATCGCCCAGCGCG | | |
| Insertion validation | FT930_ecto_1750_fw | AATTGCGTTAAATAATATCATTCA | clones | 5424 bp |
| | 253-DNlox71 | TTCACGTTACTAAAGGGAATGTA | | |
| *comX*::$P_{32}$-*cat* | | | | |
| Up recombination arm | FT694_comX_locus_fw | TGACCATGTTACACAAGCCTATATCCT | DGCC12653 chromosome | 1254 bp |
| | FT_693_comXcat_rec_rv | CGCCCTTATGGGATTTATCTTCCTTACTTCGTTTCTTTGCATAACTTCGTCTTAAT | | |
| Resistance cassette | 252-Uplox66 | TAAGGAAGATAAATCCCATA | pNZ5319 | 1402 bp |
| | 253-DNlox71 | TTCACGTTACTAAAGGGAATGTA | | |
| Down recombination arm | FT692_comXcat_rec_fw | TCTACATTCCCTTTAGTAACGTGAACCATGACCATTTTATAGGTTTAGATGTTTATG | DGCC12653 chromosome | 1505 bp |
| | 288_AR_ComX_DN_luxR | CCCAACATCTCACGACAC | | |
| Insertion validation | 252-Uplox66 | TAAGGAAGATAAATCCCATA | clones | 3343 bp |
| | CP_comXRVdiag | ATTCCTTAGAAAGGAGGTGATC | | |
| *comEC*::$P_{32}$-*cat* | | | | |
| Up recombination arm | FT685_comEC_DGCC_locus_fw | GTGAAGATCAGCCAACCACTCTTTCCA | DGCC12653 chromosome | 974 bp |
| | FT684_comEC_DGCC_rec_rv | CGCCCTTATGGGATTTATCTTCCTTAGTAAAGGCAATAAGATTACATCAAATAA | | |
| Resistance cassette | 252-Uplox66 | TAAGGAAGATAAATCCCATA | pNZ5319 | 1402 bp |
| | 253-DNlox71 | TTCACGTTACTAAAGGGAATGTA | | |
| Down recombination arm | FT 686 comEC DGCC rec fw | ATCTACATTCCCTTTAGTAACGTGAATATTAGGAACTTTTCTCTTGTCTCTAATTGG | DGCC12653 chromosome | 1227 bp |
| | FT 687 comEC DGCC locus rv | TCAAAGTGTGCGCTGTGAAGTCATTACTC | | |
| Insertion validation | 252-Uplox66 | TAAGGAAGATAAATCCCATA | clones | 3445 bp |
| | BlD_diagINTcomECRV1 | GTCCAATAATACCATTCATGAAC | | |
| *DGCC12653_00250*::*spc* | | | | |
| Up recombination arm | FT772_00250_locus_fw | ATTGATGAAGTTGGTCGTGGA | DGCC12653 chromosome | 1284 bp |
| | FT773_00250_rec_rv | ATCCTTATGGGATTTATCTTCCTTACATCAAATCTATGCGCTCAAAACTCATCAT | | |
| Resistance cassette | 252-Uplox66 | TAAGGAAGATAAATCCCATA | pJUD-*spc* | 1034 bp |
| | 253-DNlox71 | TTCACGTTACTAAAGGGAATGTA | | |
| Down recombination arm | FT778_00250_rec_fw | ATTACATTCCCTTTAGTAACGTGAATAGTCTCTTTTGAGCTGTTGCTT | DGCC12653 chromosome | 1496 bp |
| | FT779_00250_locus_rv | GTTATTGTATGACATTAAGCATC | | |
| Insertion validation | 252-Uplox66 | TAAGGAAGATAAATCCCATA | clones | 2505 bp |
| | FT780_00250_diagR | TCCGTCAGTCACAACGTCTGG | | |
| *DGCC12653_03130*::*spc* | | | | |
| | FT781_03130_locus_fw | GGAGCGTTTAGACGCACCAACA | | 1216 bp |

| | | | | |
|---|---|---|---|---|
| Up recombination arm | FT782_03130_rec_rv | ATCCTTATGGGATTTATCTTCCTTAACCATTCCACAT TCCAGCATTTCAATTTA | DGCC12653 chromosome | |
| Resistance cassette | 252-Uplox66 | TAAGGAAGATAAATCCCATA | pJUD-*spc* | 1034 bp |
| | 253-DNlox71 | TTCACGTTACTAAAGGGAATGTA | | |
| Down recombination arm | FT783_03130_rec_fw | ATTACATTCCCTTTAGTAACGTGAAATTGGGTTTGCC AGAGATGAAGGCTTATT | DGCC12653 chromosome | 1267 bp |
| | FT784_03130_locus_rv | ATTGCTTCTGGTATGATAATAAG | | |
| Insertion validation | 252-Uplox66 | TAAGGAAGATAAATCCCATA | clones | 2449 bp |
| | FT785_03130_diagR | TCAGCCGAATAGTTTGTATAAAC | | |
| *DGCC12653_04225*::*spc* | | | | |
| Up recombination arm | FT786_04225_locus_fw | GGTTTGCGATAATCATTCTTGTA | DGCC12653 chromosome | 1151 bp |
| | FT787_04225_rec_rv | ATCCTTATGGGATTTATCTTCCTTAGGTTTGACCTAA TTCAAATCTTGATAAAC | | |
| Resistance cassette | 252-Uplox66 | TAAGGAAGATAAATCCCATA | pJUD-*spc* | 1034 bp |
| | 253-DNlox71 | TTCACGTTACTAAAGGGAATGTA | | |
| Down recombination arm | FT788_04225_rec_fw | ATTACATTCCCTTTAGTAACGTGAATAGGAAAATTA TTAACATAAAAAAGTCTC | DGCC12653 chromosome | 1265 bp |
| | FT789_04225_locus_rv | CATTACTACTTCCCTACTAACGC | | |
| Insertion validation | 252-Uplox66 | TAAGGAAGATAAATCCCATA | clones | 2436 bp |
| | FT790_04225_diagR | TAAGGAAGATAAATCCCATA | | |
| *DGCC12653_05990*::*spc* | | | | |
| Up recombination arm | FT 791 05990 locus fw | TACGGTGATTCCTGACCGTATCG | DGCC12653 chromosome | 1025 bp |
| | FT 792 05990 rec rv | ATCCTTATGGGATTTATCTTCCTTATAACTACGGTCT ATTTGAAAGCATTATATG | | |
| Resistance cassette | 252-Uplox66 | TAAGGAAGATAAATCCCATA | pJUD-*spc* | 1034 bp |
| | 253-DNlox71 | TTCACGTTACTAAAGGGAATGTA | | |
| Down recombination arm | FT 793 05990 rec fw | ATTACATTCCCTTTAGTAACGTGAATGTTCATATATG ATTATTTTTGTTTATTA | DGCC12653 chromosome | 1247 bp |
| | FT 794 05990 locus rv | CACTCCTAACTGTTTCCACATC | | |
| Insertion validation | 252-Uplox66 | TAAGGAAGATAAATCCCATA | clones | 2464 bp |
| | FT 795 05990 diagR | GCTTCTGGTCGCCTTGTTACC | | |
| *DGCC12653_07535*::*spc* | | | | |
| Up recombination arm | FT796_07535_locus_fw | ATACAGGTGGGATTGAACTTTCA | DGCC12653 chromosome | 1281 bp |
| | FT797_07535_rec_rv | ATCCTTATGGGATTTATCTTCCTTAATATACCATTTA ATCTTCTTTCAATTAAC | | |
| Resistance cassette | 252-Uplox66 | TAAGGAAGATAAATCCCATA | pJUD-*spc* | 1034 bp |
| | 253-DNlox71 | TTCACGTTACTAAAGGGAATGTA | | |
| Down recombination arm | FT798_07535_rec_fw | ATTACATTCCCTTTAGTAACGTGAAGTATGATAAAA GAATACTAAGAGAA | DGCC12653 chromosome | 1228 bp |
| | FT799_07535_locus_rv | CTATCCGTATTGCGAATTGC | | |
| Insertion validation | 252-Uplox66 | TAAGGAAGATAAATCCCATA | clones | 2351 bp |
| | FT800_07535_diagR | TAGCTTGTGTCGGTTGCAAAG | | |
| *DGCC12653_08355*::*spc* | | | | |
| Up recombination arm | FT801_08355_locus_fw | CGCAAGTACCCTTTATGGTGGA | DGCC12653 chromosome | 1258 bp |
| | FT802_08355_rec_rv | ATCCTTATGGGATTTATCTTCCTTAGCTTCTCCTTGA TAAGTAAAATATTTAACA | | |
| Resistance cassette | 252-Uplox66 | TAAGGAAGATAAATCCCATA | pJUD-*spc* | 1034 bp |
| | 253-DNlox71 | TTCACGTTACTAAAGGGAATGTA | | |
| Down recombination arm | FT803_08355_rec_fw | ATTACATTCCCTTTAGTAACGTGAACCGTAGTGATA ACACTACGGCTTTTTTGT | DGCC12653 chromosome | 1227 bp |
| | FT804_08355_locus_rv | GCCCTTATTTTATCATAAATCTTGG | | |
| Insertion validation | 252-Uplox66 | TAAGGAAGATAAATCCCATA | clones | 2585 bp |
| | FT805_08355_diagR | ATCATCACGACATCAAGTGG | | |
| *DGCC12653_09325*::*spc* | | | | |

| | | | | |
|---|---|---|---|---|
| Up recombination arm | FT806_09325_locus_fw | CATGGTTAGAACTCAATAACTATG | DGCC12653 chromosome | 1224 bp |
| | FT807_09325_rec_rv | ATCCTTATGGGATTTATCTTCCTTATTGTTTCATCTA ATTATCCTTCTTTTA | | |
| Resistance cassette | 252-Uplox66 | TAAGGAAGATAAATCCCATA | pJUD-*spc* | 1034 bp |
| | 253-DNlox71 | TTCACGTTACTAAAGGGAATGTA | | |
| Down recombination arm | FT808_09325_rec_fw | ATTACATTCCCTTTAGTAACGTGAACGCCATGTTCTT AATTTAGAAGAATCA | DGCC12653 chromosome | 1228 bp |
| | FT809_09325_locus_rv | ACGTCCACCACCACATACAGTC | | |
| Insertion validation | 252-Uplox66 | TAAGGAAGATAAATCCCATA | clones | 2385 bp |
| | FT_810_09325_diagR | CCATTACAGCAATATAAACATCCA | | |
| *DGCC12653_11640*::*spc* | | | | |
| Up recombination arm | FT811_11640_locus_fw | GACAGCTTCTCATAAGTCATC | DGCC12653 chromosome | 1239 bp |
| | FT812_11640_rec_rv | ATCCTTATGGGATTTATCTTCCTTACCATAGTATTGA TTAACCATTTGTCTTGTC | | |
| Resistance cassette | 252-Uplox66 | TAAGGAAGATAAATCCCATA | pJUD-*spc* | 1034 bp |
| | 253-DNlox71 | TTCACGTTACTAAAGGGAATGTA | | |
| Down recombination arm | FT813_11640_rec_fw | ATTACATTCCCTTTAGTAACGTGAAATCGGTGCTAC AATAAAGTTAAGGAGACA | DGCC12653 chromosome | 1278 bp |
| | FT814_11640_locus_rv | CCAAGCAAAGGTCGCTAAAGTTC | | |
| Insertion validation | 252-Uplox66 | TAAGGAAGATAAATCCCATA | clones | 2380 bp |
| | FT815_11640_diagR | ATCCGAATCACCAAGATAGGCG | | |
| *DGCC12653_12925*::*spc* | | | | |
| Up recombination arm | FT821_12925_locus_fw | GCAGTTGCCATTATTCCTGGTATC | DGCC12653 chromosome | 1305 bp |
| | FT822_12925_rec_rv | ATCCTTATGGGATTTATCTTCCTTATTTGAGTCCATA AATTATCCTCATT | | |
| Resistance cassette | 252-Uplox66 | TAAGGAAGATAAATCCCATA | pJUD-*spc* | 1034 bp |
| | 253-DNlox71 | TTCACGTTACTAAAGGGAATGTA | | |
| Down recombination arm | FT823_12925_rec_fw | ATTACATTCCCTTTAGTAACGTGAATTAGGTTATGA CAAAGCGGCAAGTTATATG | DGCC12653 chromosome | 1299 bp |
| | FT824_12925_locus_rv | GAAGCATAGCATAAATTGGAAGCG | | |
| Insertion validation | 252-Uplox66 | TAAGGAAGATAAATCCCATA | clones | 2367 bp |
| | FT825_12925_diagR | CTAATCCATCTGACCCTATGACAT | | |
| *DGCC12653_13335*::*spc* | | | | |
| Up recombination arm | FT831_13335_locus_fw | 5'TCTCGATTGGTCTAATTGTTGTTG3' | DGCC12653 chromosome | 1130 bp |
| | FT832_13335_rec_rv | 5'ATCCTTATGGGATTTATCTTCCTTATTCATTTTCCC TTTGTCATATATTCCA3' | | |
| Resistance cassette | 252-Uplox66 | TAAGGAAGATAAATCCCATA | pJUD-*spc* | 1034 bp |
| | 253-DNlox71 | TTCACGTTACTAAAGGGAATGTA | | |
| Down recombination arm | FT833_13335_rec_fw | ATTACATTCCCTTTAGTAACGTGAAAATTATAGGGC TTTGGCATATATTCCA | DGCC12653 chromosome | 1229 bp |
| | FT834_13335_locus_rv | CTGAACTAACAACGAAATTAACAAAC | | |
| Insertion validation | 252-Uplox66 | TAAGGAAGATAAATCCCATA | clones | 2317 bp |
| | FT835_13335_diagR | GACACACAATTTCCGCACCAT | | |
| *DGCC12653_01140-DGCC12653_01145*::$P_{32}$- *cat* | | | | |
| Up recombination arm | FT722_01140_locus_fw | AGTCGAACCCCTGTCCAAACAC | DGCC12653 chromosome | 1171 bp |
| | FT723_01140_rec_rv | GCCCTTATGGGATTTATCTTCCTTAAGTTACTTCGTA ATCATTTTGCCGAAAATT | | |
| Resistance cassette | 252-Uplox66 | TAAGGAAGATAAATCCCATA | pNZ5319 | 1402 bp |
| | 253-DNlox71 | TTCACGTTACTAAAGGGAATGTA | | |
| Down recombination arm | FT724_01140_rec_fw | ATCTACATTCCCTTTAGTAACGTGAAAGGAACAACT TTCACCGTCAATCTTTAATCA | DGCC12653 chromosome | 1185 bp |
| | FT725_01140_locus_rv | CTGTCAGTCGAATGACAGCA | | |
| Insertion validation | 252-Uplox66 | TAAGGAAGATAAATCCCATA | clones | 2593 bp |
| | FT726_01140_diagR | TCCAAGCGAACGTGATTTGGC | | |

| | | | | |
|---|---|---|---|---|
| *DGCC12653_02295-DGCC12653_02300*::$P_{32}$-*cat* | | | | |
| Up recombination arm | FT762_02295_locus_fw | ATGGTGAGCAAGAACGTCAGGT | DGCC12653 chromosome | 1110 bp |
| | FT763_02295_rec_rv | GCCCTTATGGGATTTATCTTCCTTATTCTTGTCCATA TTATAGCATAAAAAAACGT | | |
| Resistance cassette | 252-Uplox66 | TAAGGAAGATAAATCCCATA | pNZ5319 | 1402 bp |
| | 253-DNlox71 | TTCACGTTACTAAAGGGAATGTA | | |
| Down recombination arm | FT764_02295_rec_fw | TCTACATTCCCTTTAGTAACGTGAAGAAGATTGAAC TTGATTCCAATTATCCA | DGCC12653 chromosome | 1260 bp |
| | FT765_02295_locus_rv | CTGAAACCATATATCCAAACGC | | |
| Insertion validation | 252-Uplox66 | TAAGGAAGATAAATCCCATA | clones | 2839 bp |
| | FT766_02295_diagR | CCTTCCATGTTACCCATAGCTTGAC | | |
| *DGCC12653_03165-DGCC12653_03175*::$P_{32}$-*cat* | | | | |
| Up recombination arm | FT695_03175_locus_fw | GTAGAGTTGTCTATGTTAATGATG | DGCC12653 chromosome | 1402 bp |
| | FT696_03175_rec_rv | CGCCCTTATGGGATTTATCTTCCTTAAATCAACTTTT AAGTAATCAAGTTCATAGA | | |
| Resistance cassette | 252-Uplox66 | TAAGGAAGATAAATCCCATA | pNZ5319 | 1402 bp |
| | 253-DNlox71 | TTCACGTTACTAAAGGGAATGTA | | |
| Down recombination arm | FT697_03175_rec_fw | TCTACATTCCCTTTAGTAACGTGAATACAACAGAAG TGGCTGGTAAAACAGTCAA | DGCC12653 chromosome | 1452 bp |
| | FT698_03175_locus_rv | ACGATTGAAACTTTGACTTAACTGATG | | |
| Insertion validation | 252-Uplox66 | TAAGGAAGATAAATCCCATA | clones | 2900 bp |
| | FT699_03175_diagR | CTTAGCTCATGTGAAGCATCCGAG | | |
| *DGCC12653_03735-DGCC12653_03740*::$P_{32}$-*cat* | | | | |
| Up recombination arm | FT737_03735_locus_fw | GAGATACGTTAGCCAATGGGA | DGCC12653 chromosome | 1233 bp |
| | FT738_03735_rec_rv | GCCCTTATGGGATTTATCTTCCTTAAACGAGTCGCC AAGATAGTAAAGAGTATAG | | |
| Resistance cassette | 252-Uplox66 | TAAGGAAGATAAATCCCATA | pNZ5319 | 1402 bp |
| | 253-DNlox71 | TTCACGTTACTAAAGGGAATGTA | | |
| Down recombination arm | FT739_03735_rec_fw | TCTACATTCCCTTTAGTAACGTGAAGCAACAATTTA TGCCATTCAACATCACTTA | DGCC12653 chromosome | 1284 bp |
| | FT740_03735_locus_rv | CTCCGTAGATTGAAGAACCT | | |
| Insertion validation | 252-Uplox66 | TAAGGAAGATAAATCCCATA | clones | 2846 bp |
| | FT741_03735_diagR | GCCAGTATATGTTTCAGCTAC | | |
| *DGCC12653_04255-DGCC12653_04260*::$P_{32}$-*cat* | | | | |
| Up recombination arm | FT742_04260_locus_fw | GTGAAAATATGGCAGTTACAA | DGCC12653 chromosome | 1225 bp |
| | FT743_04260_rec_rv | GCCCTTATGGGATTTATCTTCCTTAAGAGTGATAAA TCATCCTCCACTAATAA | | |
| Resistance cassette | 252-Uplox66 | TAAGGAAGATAAATCCCATA | pNZ5319 | 1402 bp |
| | 253-DNlox71 | TTCACGTTACTAAAGGGAATGTA | | |
| Down recombination arm | FT744_04260_rec_fw | TCTACATTCCCTTTAGTAACGTGAAGTGTGGATAGC ACAATTTCAAATCCTGCA | DGCC12653 chromosome | 1302 bp |
| | FT745_04260_locus_rv | ATCATAAGCACGTTGAACAA | | |
| Insertion validation | 252-Uplox66 | TAAGGAAGATAAATCCCATA | clones | 2896 bp |
| | FT746_04260_diagR | GAATCACAACTGGTTCTTCAT | | |
| *DGCC12653_04650-DGCC12653_04655*::$P_{32}$-*cat* | | | | |
| Up recombination arm | FT747_04655_locus_fw | CTCGGCACTAAATGTATCTT | DGCC12653 chromosome | 1198 bp |
| | FT748_04655_rec_rv | GCCCTTATGGGATTTATCTTCCTTAGCTTTCACAATA ATTTCATCATCTTCTAC | | |
| Resistance cassette | 252-Uplox66 | TAAGGAAGATAAATCCCATA | pNZ5319 | 1402 bp |
| | 253-DNlox71 | TTCACGTTACTAAAGGGAATGTA | | |
| Down recombination arm | FT749_04655_rec_fw | TCTACATTCCCTTTAGTAACGTGAATGAACATGGAT TTACTGGATATAATGGTC | DGCC12653 chromosome | 1282 bp |
| | FT750_04655_locus_rv | TTAAGACTAAGAAAGTCCTATCTT | | |
| | 252-Uplox66 | TAAGGAAGATAAATCCCATA | clones | 2836 bp |

| | | | | |
|---|---|---|---|---|
| Insertion validation | FT751_04655_diagR | GCCAAACATAATACTACGATGT | | |
| *DGCC12653_06415-DGCC12653_06420*::$P_{32}$-*cat* | | | | |
| Up recombination arm | FT752_06420_locus_fw | GGCAACAAGACGACAGGAACGA | DGCC12653 chromosome | 1175 bp |
| | FT753_06420_rec_rv | GCCCTTATGGGATTTATCTTCCTTATCATAGTTGTCC TTTCAAGTGGAA | | |
| Resistance cassette | 252-Uplox66 | TAAGGAAGATAAATCCCATA | pNZ5319 | 1402 bp |
| | 253-DNlox71 | TTCACGTTACTAAAGGGAATGTA | | |
| Down recombination arm | FT754_06420_rec_fw | TCTACATTCCCTTTAGTAACGTGAAGATGATGGTAC TGATGAGTTTGAATTTG | DGCC12653 chromosome | 1215 bp |
| | FT755_06420_locus_rv | CGCGGCTTCTGTTGCTGGTT | | |
| Insertion validation | 252-Uplox66 | TAAGGAAGATAAATCCCATA | clones | 2680 bp |
| | FT751_04655_diagR | GATTCAATGCTGCAACTGGTCT | | |
| *rpsL** donor DNA | | | | |
| *rpsL** locus | BlD-LLcfusARpsL | ACACCTTTGTTCTTGAAGG | IL1403 streptomycine$^r$ | 3712 bp |
| | BlD-LLldacARpsL | AAGCTGTCAGTAAATTGACG | | |
| *rpsL** sequencing | | | | |
| *rpsL** gene | BlD-RpsLUnivUp | GTATTTCTCATCGCTTCGC | IL1403 streptomycine$^r$ | 525 bp |
| | BlD-RpsLUnivDown | CCAAAACTTCACGTTTTGG | | |
| pGh$P_{comX}$-*luxAB* | | | | |
| $P_{comX}$ | FT886_PcomX_fw | AATAAAAAAGCAAGGTAAATAGCC | DGCC12653 chromosome | 179 bp |
| | FT887_PcomX_rv | GTATGTAAGCAAAAAGTTTCCAAATTTCATTGGAAA GTTCTCCTTTTATATTATC | | |
| pGh-*luxAB* | FT884_pGhPcomGAlux_fw | ATGAAATTTGGAAACTTTTTGCTTACATAC | pGh$P_{comGA[MG]}$-*luxAB* | 5808 bp |
| | FT925_PcomXlux_rec_rv | AAGGCTATTTACCTTGCTTTTTTATTCTCGAGGGGGG GCCCGGTACCCAATTCGC | | |
| pGh$P_{comGA}$-*luxAB* | | | | |
| $P_{comGA}$ | LuxcIOF1_XhoI | ATAGTCTCGAGAAATAAATGGCTACAAAATT | IO-1 chromosome | 479 bp |
| | LuxIOR1 | GTAAGCAAAAAGTTTCCAAATTTCATACTAGACTAT ACGCAAATAATC | | |
| pGh-*luxAB* | LuxcIOR2_XhoI | ATAGCTCGAGTCCCCTGACGAACTCAAGAAGATGC | pGh$P_{comGA[MG]}$-*luxAB* | 5934 bp |
| | LuxIOF2 | GATTATTTGCGTATAGTCTAGTATGAAATTTGGAAA CTTTTTGCTTAC | | |
| pGh$P_{comGA}$-*luxAB* | | | | |
| $P_{comX}$ | FT886_PcomX_fw | AATAAAAAAGCAAGGTAAATAGCC | DGCC12653 chromosome | 287 bp |
| | FT968_sfGFP_rec | CACCTGTGAACAGCTCTTCTCCTTTTGACATTGGAA AGTTCTCCTTTTATATTAT | | |
| *gfp*$^{sf}$ gene | FT967_sfGFP_fw | ATGTCAAAAGGAGAAGAGCTGTTCACAGGT | pDR111-sfGFP(Bs) | 737 bp |
| | FT969_sfGFP_rv_SacII | CTACTGTCCCGCGGTCATTATTACTTATAAAGCTCAT CCATGCCGT | | |
| pGh | FT970_pGh_SacII | ATGTCTACCGCGGATCCTCTAGAGTCCGCTAGGG | pG$^+$host9 | 3721 bp |
| | FT925_PcomXlux_rec_rv | AAGGCTATTTACCTTGCTTTTTTATTCTCGAGGGGGG GCCCGGTACCCAATTCGC | | |
| pBAD_6his-*ccpA* | | | | |
| *ccpA* | FT1447_ccpA_rec_fw | GGCATCACCATCACCATCACGTAGAATCAACAACA ACAATTTATGATGTGGCA | DGCC12653 chromosome | 1036 bp |
| | FT1448_ccpA_rec_rv | CCAAAACAGCCAAGCTTCTATTATTTGGTAGAACGA CGAGAAAAGATTTCATG | | |
| pBAD-6his | FT1328_pBAD_stop_fw | TAGAAGCTTGGCTGTTTTGGCGGATGAG | pBAD-6his | 3985 bp |
| | FT1446_pBAD6his_rev | GTGATGGTGATGGTGATGCCCCATG | | |
| pNZ8048_6his-*codY* | | | | |
| 6his-*codY* | FT1321_6HiscodY_Up | ATTATAAGGAGGCACTCACCATGGGGCATCACCATC ACCATCACGTGGCTACATTACTTGAAAAAAC | DGCC12653 chromosome | 853 bp |
| | FT1322_6HiscodY_Dw | CCAAAACAGCCAAGCTTCTATTATTTGGTAGAACGA CGAGAAAAGATTTCATG | | |
| pNZ8048 | FT1112_pNZ_Gib_fw | TGAACCAAAATTAGAAAACCAAGGCTTG | pNZ8048 | 3296 bp |
| | FT1111_pNZ_Gib_rv | GGTGAGTGCCTCCTTATAATTTATTTTG | | |
| pNZ8048_6his-*covR* | | | | |

| | | | | |
|---|---|---|---|---|
| 6his-*covR* | FT1334_pNZ_covR_rec_fw | ATTATAAGGAGGCACTCACCATGGGGCATCACCATCACCATCACACTTCAAAGAAAATTTTGATTATTG | DGCC12653 chromosome | 754 bp |
| | FT1335_pNZ_covR_rec_rv | GGTTTTCTAATTTTGGTTCATTATTTGCGTTCACGCATTACATAACCTAAGC | | |
| pNZ8048 | FT1112_pNZ_Gib_fw | TGAACCAAAATTAGAAAACCAAGGCTTG | pNZ8048 | 3296 bp |
| | FT1111_pNZ_Gib_rv | GGTGAGTGCCTCCTTATAATTTATTTTG | | |
| *mecA* locus for gene swapping | | | | |
| *mecA* | FT1319_mecA_Up_Up | TGATAAATTCCATAACAGAAACTTTGTCAT | DGCC12653 and DGCC12671 chromosome | 5030 bp |
| | FT1320_mecA_Dw_Dw | GACAAATATTAGACCTTAAAAATCCGGATG | | |
| EMSA: $P_{comX}$ | | | | |
| $P_{comX}$ | FT851_Cy3_PcomX_up | Cy3-AATAAAAAAGCAAGGTAAATAGC | DGCC12653 chromosome | 256 bp |
| | FT853_Cy5_PcomX_dw | Cy5-TGGAAAGTTCTCCTTTTATATTATC | | |
| EMSA: X2 Fw | | | | |
| $P_{comX}$ | FT888_PcomX2 | GTAATAAAAGCCATAAAAATGTCTTG | DGCC12653 chromosome | 154 bp |
| | FT853_Cy5_PcomX_dw | Cy5-TGGAAAGTTCTCCTTTTATATTATC | | |
| EMSA: X3 Fw | | | | |
| $P_{comX}$ | FT1558 | TGTCTTGGAAGGATAGATATGAAAATAGCTCATTATAAT | DGCC12653 chromosome | 135 bp |
| | FT853_Cy5_PcomX_dw | Cy5-TGGAAAGTTCTCCTTTTATATTATC | | |
| EMSA: X4 Fw | | | | |
| $P_{comX}$ | FT889 | AAGATGCTCCCTGACTGTTTATTAGAT | DGCC12653 chromosome | 96 bp |
| | FT853_Cy5_PcomX_dw | Cy5-TGGAAAGTTCTCCTTTTATATTATC | | |
| EMSA: X5 Rv | | | | |
| $P_{comX}$ | FT851_Cy3_PcomX_up | Cy3-AATAAAAAAGCAAGGTAAATAGC | DGCC12653 chromosome | 86 bp |
| | FT1355 | CGTGTTTTACCTAACGATATTA | | |
| EMSA: X6 Rv | | | | |
| $P_{comX}$ | FT851_Cy3_PcomX_up | Cy3-AATAAAAAAGCAAGGTAAATAGC | DGCC12653 chromosome | 115 bp |
| | FT975 | TGGCTTTTATTACATTATAATAATTTTTACGT | | |
| EMSA: $P_{comX}$ native CodY-box | | | | |
| $P_{comX}$ | FT1368_PcomX_codY_control | TATTTATTGGGTTATAAATTCTGAATATTCGCATAATATCGTTAGGTAAAAC | DGCC12653 chromosome | 225 bp |
| | FT853_Cy5_PcomX_dw | Cy5-TGGAAAGTTCTCCTTTTATATTATC | | |
| EMSA: $P_{comX}$ mutated CodY-box | | | | |
| $P_{comX}$ | FT1369_PcomX_codY_mute | TATTTATTGGGTTATAAATTATAAATATTCGCATAATATCGTTAGGTAAAAC | DGCC12653 chromosome | 225 bp |
| | FT853_Cy5_PcomX_dw | Cy5-TGGAAAGTTCTCCTTTTATATTATC | | |
| EMSA: negative control | | | | |
| CDS *dnaE* | AK350 | CCTGTAGTTCCCTTACATAC | *S. salivarius* HSISS4 chromosome | 150 bp |
| | AK303 | TTCCATTTCTTGAGGCGAG | | |
